# Methods for evaluating bacterial dispersal on hyphal networks

**DOI:** 10.64898/2026.04.17.719220

**Authors:** Aureline Bouchard, Martin Darino, Guillaume Cailleau, Achala Narayanan, Melissa Cravero, Peter G. Kennedy, Buck T. Hanson, Aaron Robinson, Julia M. Kelliher, Kim E. Hammond-Kosack, Patrick S.G. Chain, Saskia Bindschedler, Pilar Junier

## Abstract

In heterogeneous environments, the hyphae of filamentous fungi and oomycetes can facilitate the dispersal of other microorganisms. The use of these “fungal highways” (FH) is regulated by both physical and biological factors with their interplay resulting in variable capabilities of different microbes to establish FH. Several devices have been developed to test the movement of bacteria across mycelium. However, these methods are usually time-consuming and cannot be applied either at a large scale or in a high throughput format. In this study, we developed 3D-printed experimental devices that physically separate two environments while allowing hyphal networks to act as bridges for bacterial movement. The final design allows for the simultaneous testing of up to 10 pairs and the inclusion of any culturing media. With these devices, we investigated how fungal–bacterial pairing, nutrient conditions, and inoculation strategies influence FH formation. Bacterial transport was limited in nutrient-rich media but increased under poorer nutrient conditions, consistent with enhanced exploratory growth of the mycelium. Both cis- and trans-inoculation supported FH formation, although bacterial arrival was delayed in the absence of co-inoculation. The devices were used to demonstrate that transport of bacteria by FH was relevant for the colonization of a natural substrate. Finally, we established a novel *in planta* assay to evaluate FH formation during host colonization. This assay demonstrated that *Fusarium graminearum* can transport bacteria during wheat spike colonization. Together, these results provide accessible, scalable tools to study hyphal-mediated bacterial dispersal and highlight the combined role of biological specificity and nutrient context in the establishment of FH.

**Graphical abstract:** 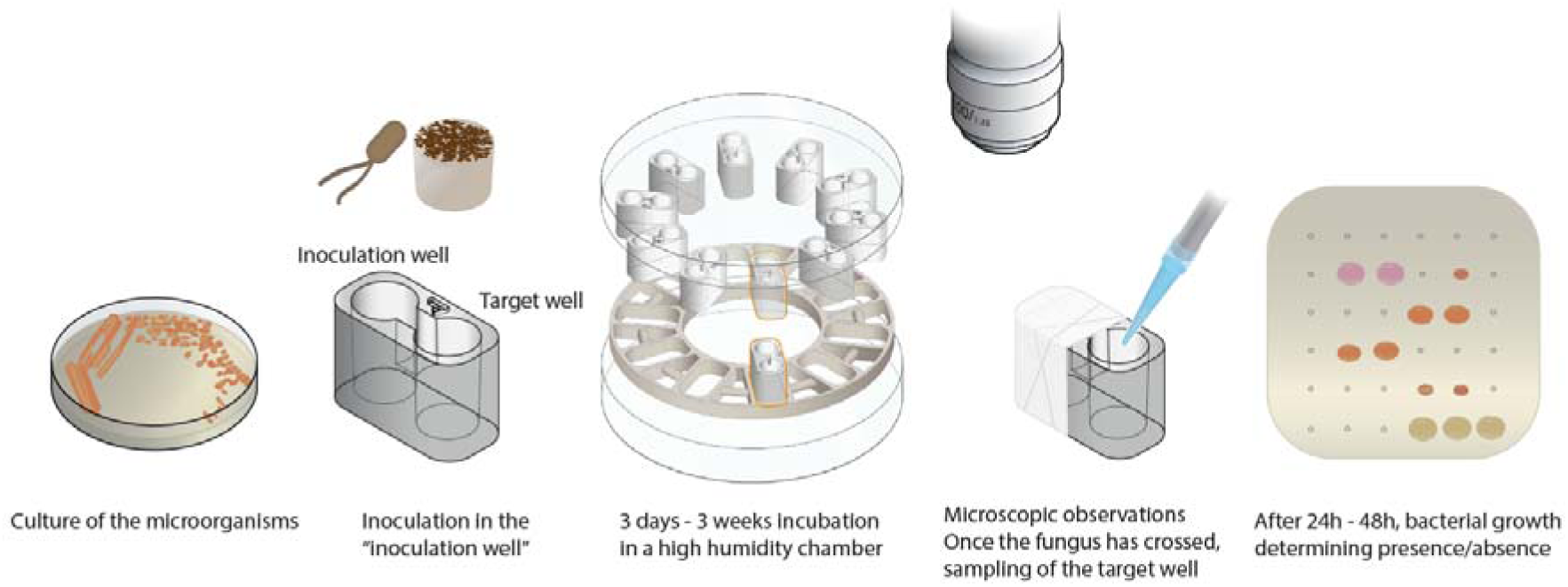

## Introduction

Most ecosystems are characterized by frequent microbial interactions that span across species and kingdom boundaries. Bacteria and fungi are crucial members of these microbial associations. At the level of individual cells or discrete populations, the effect of interactions can be seen in changes in the growth, physiology, reproduction, stress resistance, or even pathogenicity of the involved partners (Deveau et al., 2018). These individual-scale effects can lead to emergent phenotypes that have a significant impact all the way up to ecosystem functioning (Deveau et al., 2018; Frey-Klett et al., 2011; Zhou et al., 2022). For instance, the dispersal of microbial species on the mycelium of filamentous microorganisms that occurs in heterogenous environments, such as soils, can lead to shifts in community structure and dynamics due to species redistribution (Deveau et al., 2018).

The multicellular network structure that constitutes the mycelium of both filamentous fungi and oomycetes is well suited for colonization of spatially structured environments such as soils (Fischer & Glass, 2019; Fricker et al., 2007; Harris, 2008). These mycelial networks can be exploited by unicellular microorganisms, including bacteria (Furuno et al., 2010; Kohlmeier et al., 2005) and bacterial viruses (Périat et al., 2024; You, Kallies, et al., 2022; You, Klose, et al., 2022) as a scaffold for their active or passive dispersal between discontinuous environmental patches. This type of interaction is known under the general term of fungal highways (FH), regardless of whether the scaffold-forming filamentous microorganism is a true fungus or a filamentous oomycete (Furuno et al., 2010; Pion et al., 2013).

Various species of fungi, oomycetes, and bacteria have been shown to interact through FH with diverse resulting outcomes. Mutual benefits from the interaction have been demonstrated, for instance, by the exchange of key metabolites (Lohberger et al., 2019) or by the improved degradation of specific substrates (Wick et al., 2007). In the context of FH, the ability to modify traits such as morphology, motility, stress resistance or surface properties plays a significant role in the success of the interaction (Buffi et al., 2025; Fischer & Glass, 2019; Jia et al., 2024; Jiang et al., 2021). However, several FH studies to date are based on the artificial pairing of species isolated separately. Therefore, to gain a more unbiased overview of the diversity of FH partners in natural ecosystems, several experimental devices have been developed over the years to be able to “bait” FH pairs directly from environmental samples (Junier et al., 2021; Simon et al., 2015, 2017). Nevertheless, upon isolation, FH interactions still must be validated experimentally.

Numerous devices with different levels of technical complexity have been proposed to conduct this experimental validation (Bravo et al., 2013; Buffi et al., 2023; Gimeno et al., 2021). In most cases, the fundamental principle of the validation remains the same: demonstrating co-colonization of two physically separated environments. Some of the methods, such as the use of unsaturated microfluidics (Clark et al., 2024; Richter et al., 2022) or fungal drops (Buffi et al., 2023) enable microscopic observations during dispersal. Alternatively, in the split plate method, co-colonization can be demonstrated by regrowth or specific phenotypic observations such as degradation of a substrate (Bravo et al., 2013) or tracking of a fluorescently labelled bacterium (Junier et al., 2021). A commonality to existing systems is the limited throughput of the testing method. This represents a considerable limitation for the assessment of FH in large co-culture collections or when varying nutritional or physicochemical conditions. As such, increasing the throughput while retaining the potential to vary the type and composition of the growth medium represents a strong motivation for development of novel methods.

Building on our previous experience with the design and production of devices generated by additive printing (Junier et al., 2021; Kuhn et al., 2022), we developed a new device for testing FH. Three-dimensional (3D) printing offers a fast translation between digital design and finished product, allowing the testing of intricate designs with greater reproducibility, and faster production times (Iftekar et al., 2023). Accordingly, the device design was based on fast conception, prototyping, evaluation, and re-conception cycles until an optimal device topology was defined. This optimal topology was then used to evaluate the effect of various nutritive media, inoculation conditions, and mycelial-forming partners on the effectiveness of bacterial transport. For this, multiple microbial species were compared using different concentrations of the same nutrient media. Next, the inoculation scheme (i.e., co-inoculation or inoculation in separate wells) and the creation of simple nutrient gradients (i.e., dilution of the nutrient medium) were evaluated with a restricted number of bacterial-fungal pairings. Based on the results, two specific experiments were performed to evaluate the ecological relevance of the observed FH. First, non-model organisms were tested when growing on fungal necromass as a natural substrate to evaluate the potential effect of FH on carbon decomposition. Necromass was chosen because it is an abundant, yet fast cycling pool of soil organic matter that varies considerably in chemical composition (Maillard et al., 2023; Ryan et al., 2020; Wang et al., 2021). This is of particular interest because individual bacteria seem to be much less active necromass decomposers than fungi in spatially structured environments (Pérez-Pazos et al., 2024) but not in liquid settings (Narayanan et al. 2025). This suggested the potential importance of fungal physical structures in soil environments. Specifically, melanin, a recalcitrant aromatic polymer present in many fungal cell walls, interacts with nitrogen-rich fungal cell wall components to reduce carbon and nitrogen bioavailability during decomposition (Beidler et al., 2025; Maillard et al., 2023). Thus, using necromass with varying levels of melanin provides model substrate to test how microbes associate in FH with differing substrate complexity (Beidler et al., 2020; Fernandez & Kennedy, 2018; Maillard et al., 2023). Finally, *in planta* experiments were done to validate the observations made in the devices in the context of host colonization. For this *Fusarium graminearum*, a fast-growing Ascomycota of major economic importance, was chosen for its known ability to associate with bacteria and enhance virulence of the fungus (Ali et al., 2022).

## Material and methods

### Design and printing of the devices

The development of the devices was inspired by the bacterial bridge device described previously to evaluate the movement of motile bacteria across two separate containers (Kuhn et al., 2022). For printing, two types of resin were used. Initial prototyping in which the goal was to validate the printing quality, was performed with a Formlabs White resin V4, which has a lower cost. This resin was selected for printing the holder device (see results). When the design was validated, the certified biocompatible BioMed Clear resin V1 was used to print devices for microbiological experiments. This biocompatible material is suitable for common sterilization methods and was used for all the 2-wells device designs. Both resins support a printing resolution of up to 50 µm.

The various designs were created using Fusion 360 (Autodesk, San Francisco, CA 94105 USA) and saved as .stl files (**Supplementary information 1**). These files were then imported into PreForm® (Formlabs, Somerville, MA 02143 USA) to prepare the models for 3D printing on a Formlabs Form 3B+ stereolithography (SLA) printer. Following the manufacturer’s recommendations, rafts and supports were used during printing to prevent deformation. After printing, the devices were cleaned by rinsing them repeatedly in 90% isopropanol and cured using UV light and heat during 60 min at 60°C, as specified by the resin manufacturer.

### Media used

The media used in this study are described in **Supplementary Table 1**. All the media were sterilized by autoclaving at 121°C for 20 min. All the media were stored at room temperature, except those containing antibiotic or fungicide, which were kept refrigerated.

### Necromass preparation

For the preparation of the necromass medium, the fungus *Hyaloscypha bicolor*, a mycorrhizal fungus broadly distributed among northern latitude forests (Grelet et al., 2009; Kohout et al., 2011) was selected. *H. bicolor* exhibits two distinct melanin phenotypes that are easily manipulated in the lab and thus make this species a useful experimental model (Maillard et al., 2023). Varied oxygen availability and shaking speed between the two phenotypes results in a 4-fold difference in melanin content and can be visually confirmed (Fernandez & Kennedy, 2018). We prepared *H. bicolor* according to this method. In brief, *H. bicolor* was grown on cellulose membranes on 50% PDA plates, and after 3 weeks plugs from the culture were transferred to 125 mL flasks filled with either 40 mL (low melanin) or 80[mL (high melanin) 50% PDB, and shaken at 150 or 250 rpm, respectively. After one month of growth, residual media was washed off, and the biomass was freeze-dried and ground before use. To prepare necromass for inoculation into the bridges, five g of each necromass phenotype was dissolved in one L of basal media **(Supplementary Table 1**).

### Choice of fungal and bacterial strains and culture conditions

To evaluate the applicability of our method across diverse mycelial-forming microorganisms, we selected organisms representing a broad range of phylogenetic and phenotypic traits. For fungi (**Table 1**), we included *Coprinopsis cinerea*, a slow-growing Basidiomycota widely used in mycology research and previously shown to support bacterial movement along hyphae (Stajich et al., 2010; Junier et al., 2021). *Fusarium graminearum*, a fast-growing phytopathogen Ascomycota (Ali et al., 2022). We also included *Trichoderma rossicum*, a fast-growing soil fungus commonly found in decaying wood, making it ecologically relevant (Tang et al., 2022). The strain selected was shown to establish FH in previous studies (Bravo et al., 2013). To broaden the phylogenetic scope, we added *Pythium ultimum*, an oomycete previously demonstrated to transport bacteria (Furuno et al., 2012).

**Table 1.**
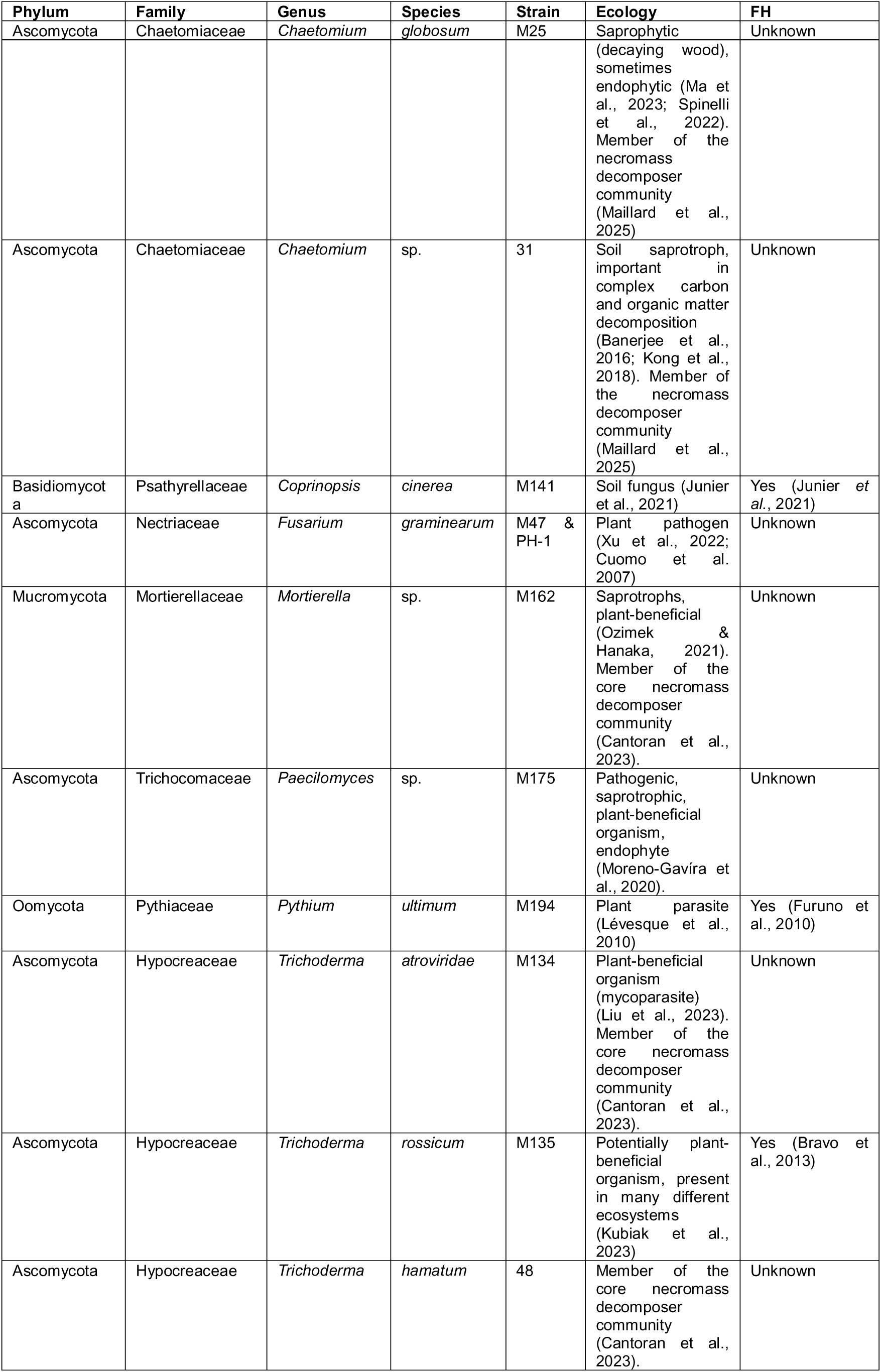
Fungi and oomycete used in this study. The organisms are organized by alphabetic order based on genera. Ecology was obtained based on a literature search. FH capability is also based on previous publications.

As bacterial partners, we selected strains with contrasting behaviors (**Table 2**). *Escherichia coli* K-12 served as a negative control, as a prior study showed it cannot disperse on fungal hyphae (Pion et al., 2013); here, we used a GFP-tagged derivative (5kfu GFP) for easy visualization of FH transport. Two *Pseudomonas koreensis* strains isolated from the *Morchella* bacteriome were included due to their established fungal associations (Cailleau et al., 2023), while *Pseudomonas putida* acted as a positive control for hyphal dispersal (Pion et al., 2013). Finally, *Paraburkholderia hospita* was chosen based on evidence that close relatives, such as *Paraburkholderia terrae*, efficiently move along fungal networks (Nazir et al., 2014; Pratama et al., 2020; Warmink et al., 2011).

**Table 2.**
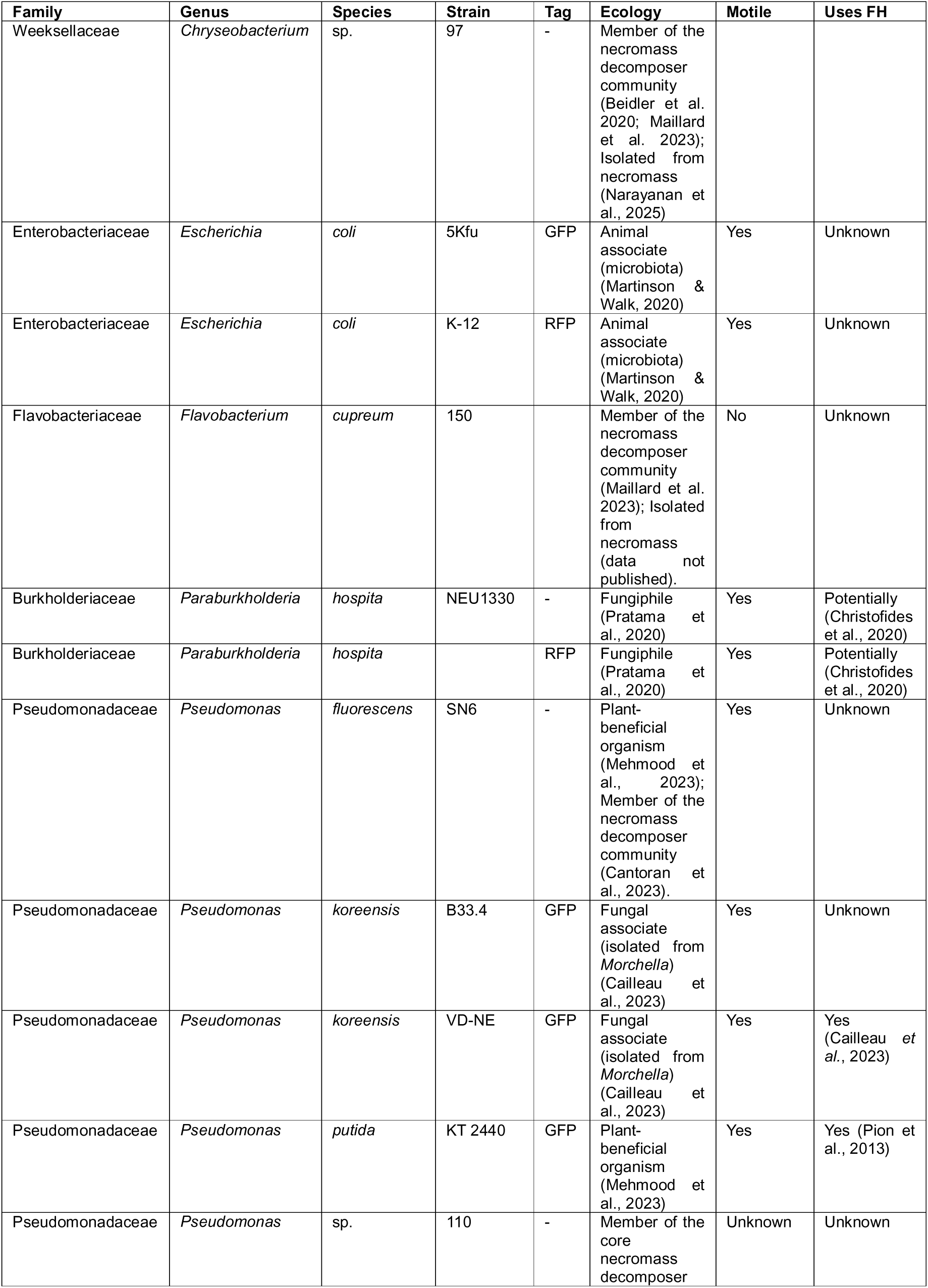
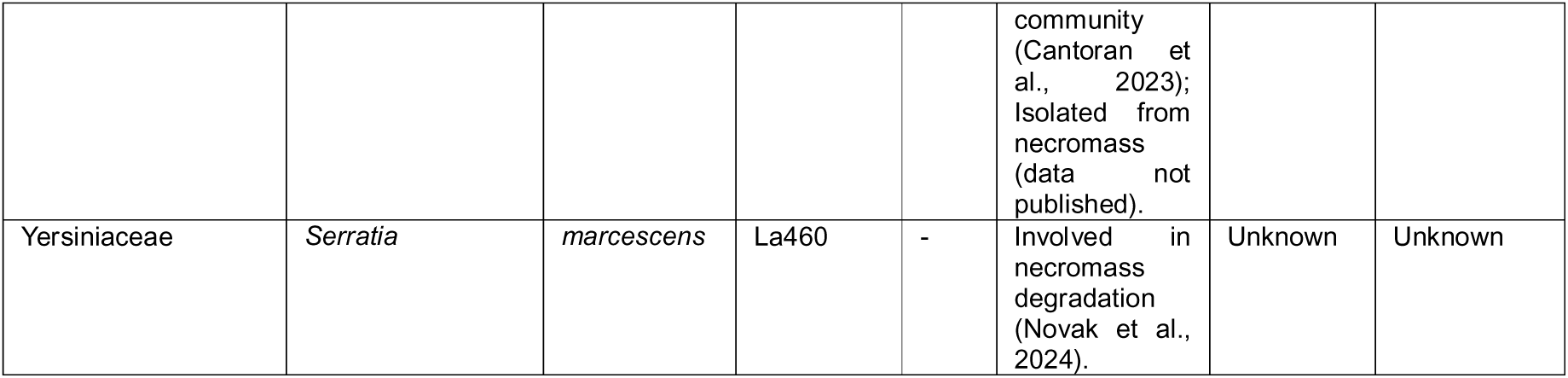
Bacterial strains used in this study. The “Tag” column indicates strains with fluorescent tags when applicable. GFP= green fluorescent protein. Ecology and use of FH (Uses FH) is based on previous literature search.

| Family | Genus | Species | Strain | Tag | Ecology | Motile | Uses FH |
| --- | --- | --- | --- | --- | --- | --- | --- |
| Weeksellaceae | *Chryseobacterium* | sp. | 97 | - | Member of the necromass decomposer community (Beidler et al. 2020; Maillard et al. 2023); Isolated from necromass (Narayanan et al., 2025) |  |  |
| Enterobacteriaceae | *Escherichia* | *coli* | 5Kfu | GFP | Animal associate (microbiota) (Martinson & Walk, 2020) | Yes | Unknown |
| Enterobacteriaceae | *Escherichia* | *coli* | K-12 | RFP | Animal associate (microbiota) (Martinson & Walk, 2020) | Yes | Unknown |
| Flavobacteriaceae | *Flavobacterium* | *cupreum* | 150 |  | Member of the necromass decomposer community (Maillard et al. 2023); Isolated from necromass (data not published). | No | Unknown |
| Burkholderiaceae | *Paraburkholderia* | *hospita* | NEU1330 | - | Fungiphile (Pratama et al., 2020) | Yes | Potentially (Christofides et al., 2020) |
| Burkholderiaceae | *Paraburkholderia* | *hospita* |  | RFP | Fungiphile (Pratama et al., 2020) | Yes | Potentially (Christofides et al., 2020) |
| Pseudomonadaceae | *Pseudomonas* | *fluorescens* | SN6 | - | Plant-beneficial organism (Mehmood et al., 2023); Member of the necromass decomposer community (Cantor et al., 2023). | Yes | Unknown |
| Pseudomonadaceae | *Pseudomonas* | *koreensis* | B33.4 | GFP | Fungal associate (isolated from *Morchella*) (Cailleau et al., 2023) | Yes | Unknown |
| Pseudomonadaceae | *Pseudomonas* | *koreensis* | VD-NE | GFP | Fungal associate (isolated from *Morchella*) (Cailleau et al., 2023) | Yes | Yes (Cailleau et al., 2023) |
| Pseudomonadaceae | *Pseudomonas* | *putida* | KT 2440 | GFP | Plant-beneficial organism (Mehmood et al., 2023) | Yes | Yes (Pion et al., 2013) |
| Pseudomonadaceae | *Pseudomonas* | sp. | 110 | - | Member of the core necromass decomposer community (Cantor et al., 2023); Isolated from necromass (data not published). | Unknown | Unknown |
| Yersiniaceae | *Serratia* | *marcescens* | La460 | - | Involved in necromass degradation (Novak et al., 2024). | Unknown | Unknown |

In addition to those organisms with previous FH knowledge, we also included a subset of microorganisms co-isolated as part of research identifying decomposers of necromass (**Tables 1 and 2**). Specifically, we used *Mortierella* sp., *Paecilomyces* sp., *Trichoderma atroviride,* and *Chaetomium globosum*. *Mortierella* and *Trichoderma* are established members of the core fungal necromass community (Cantoran et al., 2023) and available in our culture collection. Additionally, we used two strains (*Mortierella* sp. 81 and *Trichoderma hamatum* sp. 48) that were isolated from *Hyaloscypha bicolor* fungal necromass decomposed for one month in a *Pinus strobus* dominated stand at the Cedar Creek Ecosystem Science Reserve in central Minnesota, USA (see Maillard et al. 2025 and Narayanan et al. 2025 for specific details on necromass incubation and culture isolation).

Furthermore, we chose *Pseudomonas, Flavobacterium cupreum,* and *Chryseobacterium* species due to their presence in the necrobiome community (Beidler et al., 2020; Maillard et al., 2023) and the benefit they derive from fungi when grown together on necromass (K. Beidler, personal communication). *Serratia marcescens* was selected based on its ability to grow on necromass (Novak et al., 2024) and reported migration in fungal hyphae (Tal et al., 2016). All bacteria were motile except for *F. cupreum* (**Table 2**). These additional bacterial strains (except for *Serratia marcescens* and *Pseudomonas fluorescens)* were isolated in the field from *H. bicolor* necromass at the same site in Minnesota.

Fungal strains were retrieved from the fungal cryobank and cultivated on malt agar (MA; **Supplementary Table 1**) poured as a thick layer (6 mm) into 90 mm diameter Petri dishes. The thickness of the layer was important for the transfer into the inoculation wells (see "**Set-up and inoculation with optimized design"**). The cultures were incubated at 21°C in the dark for 3 to7 days before being used in the experiments. For the *in planta* experiments (see "***In planta* validation experiments**" below), conidia from *F. graminearum* PH-1 strain (Cuomo et al., 2007) were produced following the method described previously (Darino et al., 2024). Briefly, fungal spores were plated in the centre of a synthetic nutrient agar (SNA) Petri dish (**Supplementary Table 1)**. The fungus was grown for five days under constant UV and white light at room temperature. Spore production was then induced by adding two mL of TB3 media (**Supplementary Table 1)**. Two days after spore induction, the spores were collected in sterile water and store at -80[C until use.

The seven bacterial strains from the strain collection of the Laboratory of Microbiology of the University of Neuchâtel (**Table 2**) were retrieved from the bacterial cryobank and cultivated on MA. The necromass bacterial strains were sent from the US to Switzerland on 10% tryptic soy agar (TSA) (bacteria) or potato dextrose agar (PDA) (fungi) and replated on MA. Cultures were incubated at 21°C for one to two days before being used in the experiments.

### Set-up and inoculation with optimized design

The 3D-printed holder and the 3D-printed devices were immersed in deionized water and autoclaved for 20 min at 121°C. This procedure prevents breaking during sterilization. Water from the previous step was drained inside a sterile hood. The devices were subsequently dried by placing them in a 60°C oven for multiple days within a sterile container. Before the experiment, 30 x 12 x 0.1 mm PTFE tape (Kirchhoff, Deutschland) pieces were immersed in 70% ethanol for at least 15 min and dried in a sterile hood. With sterile tweezers, a device holder for 10 bridges was placed in an empty 92 mm-diameter Petri dish.

A sterile stainless-steel straw (5 mm in diameter and 6 mm in height) was used to cut agar plugs containing actively growing mycelium. Each plug was transferred into the inoculation well (usually placed as the inner well in the holder, see results) using a sterile platinum loop. When indicated, to prevent desiccation and promote colonization, 50 µL of liquid medium (same composition as the solid medium) was added to the fungal well prior to adding the bacterial inoculum. The outer wells [hereafter referred to as "Target medium" (TM)] were filled with 100 µL of liquid medium, which consisted of either 10% malt broth (MB) or 100% MB.

A bacterial suspension from a fresh culture was prepared in physiological water (0.9% NaCl) and adjusted to an OD_600_ of 0.2. For assays in which the fungus and bacterium were inoculated in the same well, 4 µL of the bacterial suspension was pipetted directly on top of the fungal agar plug in the inoculation well. For assays in which the fungus and bacterium were inoculated in separate wells, 4 µL of bacterial suspension were added to the liquid medium in the inoculation well, while the agar plug containing the fungus was placed in the target well. The wells containing the fungal inoculum were sealed manually using PTFE tape (using sterile gloves). The Petri dishes were then closed and sealed with Parafilm. Incubation (from a few days to up to 3 weeks) was performed in a desiccator (100% humidity) at 21°C.

### Evaluation of the bacterial crossing ability

To assess bacterial crossing of the wells, the TM was sampled at defined times (a few days to 2-3 weeks post inoculation). The medium in the TM was homogenized by pipetting several times up-and-down. Then, 5 µL of liquid were sampled and deposited on nutrient agar (NA) containing 50 ppm cycloheximide (fungicide). The plate was incubated overnight at 21°C. The presence of a growing colony indicated that the TM was colonized by bacteria. For the GFP-tagged strains, fluorescence was checked under a stereoscope (Nikon SMZ18 Stereomicroscope) with a UV light source.

### Molecular identification of bacteria

A colony PCR was performed on fresh bacterial colonies when a change in the identity of the crossing bacterium was suspected (i.e., non-expected macromorphological feature). For this, a colony was transferred to a PCR tube containing 20 µl nuclease-free water. The samples were heated at 98°C for 10 min to lyse the cells. Afterwards, 1 μl of the lysate was added to 24 μl of PCR mix [2.5 μl 10x Standard Taq Reaction Buffer (New England Biolabs), 0.5 μl 10 mM dNTPs, 0.5 μl 10mM forward primer, 0.5 μl 10mM reverse primer, 0.125 μl Taq DNA polymerase (5,000 units/ml) (New England Biolabs, M0273S), 19.855 μl nuclease-free water]. The primers EUB 9-27f (3’ – GAG TTT GAT CCT GGC TCA G – 5’) and 907r (5’ – CCG TCA ATT CCT TTG AGT – 3’) were used to amplify the bacterial 16S rRNA gene (Muyzer et al., 1995). The PCR program was as follows: initial denaturation at 95°C for 1 min, followed by 40 cycles at 95°C for 15 s, 59°C for 15 s, 72°C for 30 s, completed by a final extension at 72°C for 2 min. After verification of the amplification by electrophoresis (30 min, 100V, 1.2% agarose gel), the PCR products were purified on a 96-well plate MultiScreen-PCR_µ96_ (Millipore) and quantified by fluorescence (Qubit, Invitrogen). The PCR products were then prepared as instructed by Fasteris SA [4.5 µl of the PCR product (2 – 40 ng/ μl) with 0.5 μl of either primer F or R] and sent to Genesupport (Geneva, Switzerland) for Sanger sequencing.

### Experimental Design and Statistical analyses

After validation of the final design, to determine the transport capacity experiments were run in triplicates for each mycelia-forming organism-bacterium pair. The same bacteria were placed next to each other. Only one FH-transporter was included in each Petri dish. A variable number of controls with the FH-transporter alone were included in each Petri dish. Schemes of the experimental design are provided for each experiment in the corresponding figure to facilitate understanding. Statistical analyses were done using R and RStudio (version 4.3.2). For tables and figures, the packages dplyr, kableExtra, tidyverse, pagedown, xfun and ggplot2 were used.

### *In planta* validation experiments

To evaluate the relevance of the FH pairings investigated in vitro, a new experimental protocol was developed to assess FH during host colonization using a known wheat fungal phytopathogen: *F. graminearum* PH-1 strain (Cuomo et al., 2007) paired with two bacterial species. To facilitated visualization the bacteria were tagged with fluorescent labels. For this, *E. coli* K-12 and *P. hospita* were transformed with the pBBR-RFP plasmid which constitutively expresses the red fluorescence protein (RFP) (Uzelac et al., 2017), by electroporation (Mosquito et al., 2020). Briefly, overnight bacterial cultures were adjusted to an OD_600_ of 0.8. A 10 mL aliquot was placed on ice for 10 min and then centrifuged at 3,900 rpm for 10 min at 4°C. The pellet was washed twice with 10 mL of 0.3 M sucrose at 4°C. Cells were incubated on ice for 10 min between each wash. Finally, the pellet was resuspended in 200 µL of 0.3 M sucrose. Fifty microliter aliquots of the cell suspension were electroporated with five µL of pBBR-RFP plasmid using a MicroPulser electroporator (Bio-Rad). Transformants were recovered in nutrient broth (NB) medium without antibiotic for three hours at 28°C and plated on NA plates containing 25 µg/mL gentamicin (Melford, UK). Transformant colonies were visible after 48 h at 28°C. The *E. coli* and *P. hospita* strains expressing RFP were denoted as *E. coli*-RFP and *P.hospita*-RFP, respectively.

### Bacterial and fungal co-inoculation in wheat spikes

Plants of the susceptible wheat cultivar Apogee were grown as previously described (Darino et al., 2024). Plants at the flowering stage were inoculated following an adaption of the point inoculation method (Wood et al., 2020). Wheat plants at the anthesis stage were selected for inoculation. The 9^th^ and 10^th^ spikelets of the primary wheat spike were each inoculated with 5 µL of a water droplet suspension containing 5 x10^4^ spores/mL of the *F. graminearum* PH-1 strain. The droplet was placed in the space between the palea and lemma using a 10 µl micropipette (Gilson, US). Plants were kept in a saturated humid chamber under dark conditions. After 24 hours, the plants were removed from the chamber, and the spikelets previously inoculated with the fungus were subsequently inoculated with 5 µl of either *E. coli*-RFP or *P. hospita*-RFP suspension adjusted to OD_600_=0.2. Spikes were then detached from the plants and placed upright in a 200 µl pipette-tip box filled with water. A non-inoculated wheat spike was positioned in front of each inoculated spike to increase the amount of tissue surface available for mycelia colonisation. The boxes containing the inoculated spikes were incubated in a chamber under saturated humid conditions for 5 days. As a control, spikes inoculated only with the PH-1 strain, *E. coli-RFP*, or *P. hospita*-RFP were included. Three spikes were inoculated for each treatment.

Bacterial strains for co-inoculation with the fungus were prepared as follows: *E. coli*-RFP and *P. hospita*-RFP were grown overnight in NB containing 25 µg/mL gentamicin (Melford, UK) with agitation at 28 °C. The next day, fresh NB cultures were inoculated with each overnight culture and incubated for an additional 5 hours under the same conditions. Bacterial cultures were then centrifuged at 3,500 rpm for 10 min at RT. The resulting pellet was resuspended in 10 mL of sterile water and centrifuged again under the same conditions. Finally, the pellet was resuspended in 5 mL water. For each bacterial strain, a suspension with an OD_600_=0.2 was prepared.

To evaluate bacterial dispersion beyond the inoculation site, spikelets located below the point of fungal-bacterial inoculation were collected separately from the co-inoculated spikelet with the fungus and the bacterial strains. Each sample was individually macerated using a mortar and pestle with 4 mL of sterile water, and the resulting suspensions were transferred to 15 mL Falcon tubes. Four hundred microliters of each suspension were plated onto NA plates containing 25 µg/mL gentamicin and 250 µg/mL cycloheximide (Melford, UK). In the case of the spikelets co-inoculated with the fungus and the bacterial strain, the macerated tissue suspension was serially diluted 10-fold, and 5 µL of each dilution was plated onto agar plates containing gentamicin and cycloheximide. Plates were incubated for 48h at 28°C. Around 6 colonies from each plate were transferred to fresh NA plate supplemented with 25 µg/mL gentamicin. To confirm the isolated colonies corresponded to either *E. coli* – RFP or *P. hospita* – RFP, colony PCR was performed. Briefly, a small portion of each bacterial colony was resuspended in 80 µL of sterile water, and 1 µL of this suspension was used as template to amplify the *RFP* gene using the Dream Taq DNA polymerase kit (Fisher Scientific, US), following the manufacturer’s instructions. Primer combination RFP_F1 (TTACCAAAGGTGGTCCGCTG) and RFP_R1 (TTAAGCACCGGTGGAGTGAC) amplifies a product of 536 bp from the *RFP* gene.

## Results

### Development and optimization of the 3D-printed devices

Iterative designs were evaluated to produce a device to demonstrate the establishment of FH in pairings of bacteria and mycelia-forming microorganisms. The essential feature of the device was to create two separated environments that must be connected by the mycelium to allow the colonization of the transported bacteria (**Supplementary Figure 1 and Figure 1**). Another feature of the device was the flexible use of multiple culturing media and the type of substrates (i.e., concentration of nutrients and liquid vs. solid media), as considered in previous studies (Junier et al., 2021; Simon et al., 2017).

**Figure 1.**
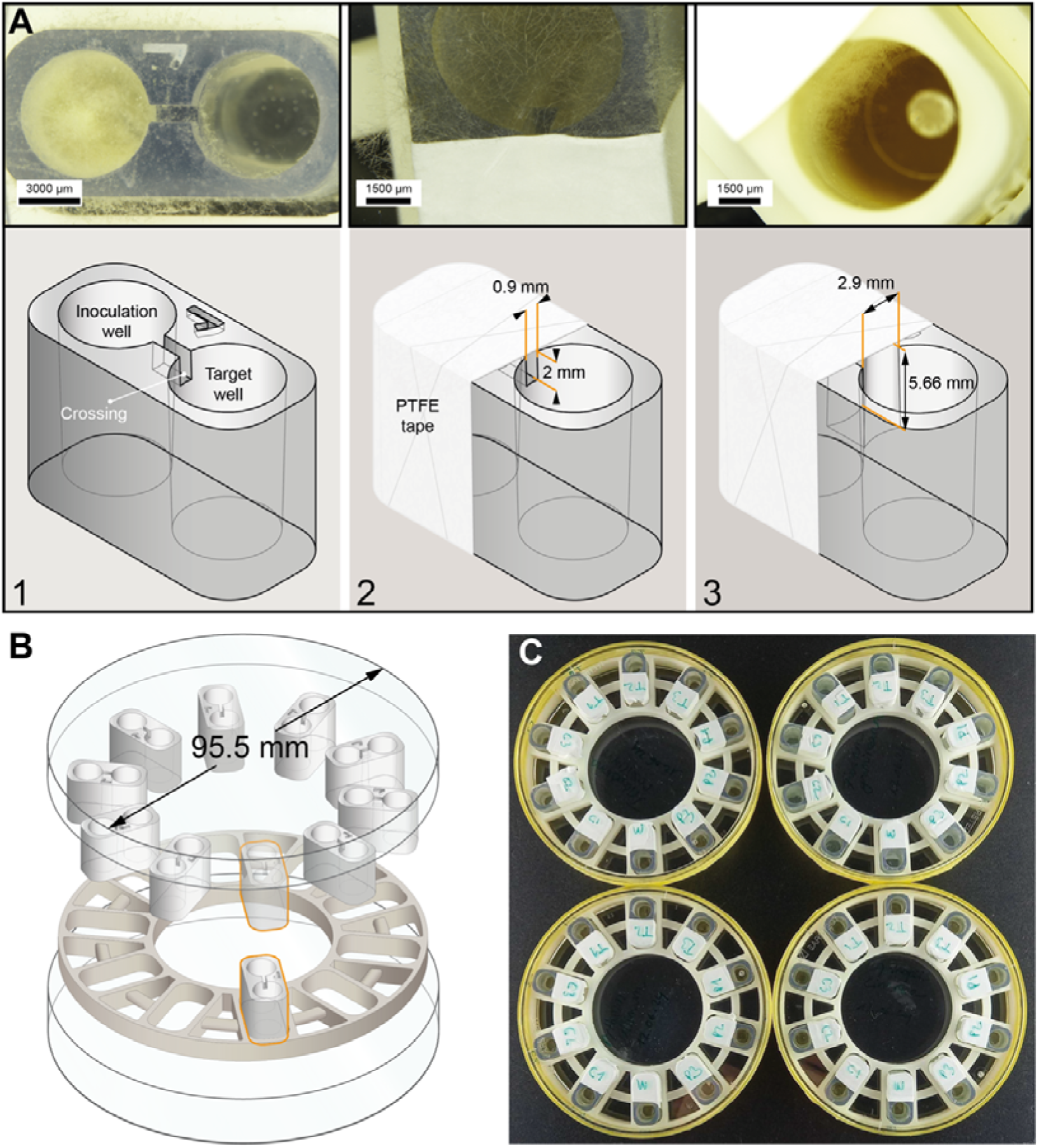
Evolution of the design of the 2-well device. (**A**) The evolution of the design led to the generation of a 2-well device based on the size of the wells of a 96-well plate. The two wells in this case were connected by a canal. The initial evaluation (1) resulted in growth beyond the device. In the next iteration, we covered the inoculation well with PTFE tape to create a hydrophobic surface reducing mycelial spread (2). However, this modification was not sufficient to reduce mycelial growth beyond the device. Finally, by using Teflon tape to cover the inoculation well and enlarging the size of the canal and making it closer to the inoculum, mycelial growth could be controlled and restricted to the interior of the device (3). (**B**) In addition to the device, we created a holder for Petri dishes. Each device can be placed in the holder to perform up to 10 simultaneous experiments at once. (**C**) Image showing the testing of the device on a real-size experiment.

To achieve higher throughput, the devices were initially dimensioned to fit between two wells of a Costar® 96-well cell culture multiplate from Corning (**Supplementary Figure 1A**). The design features of the initial device tested included: (1) beveled bridge piers for easy insertion into an inoculation well in which an agar plug (or liquid medium) containing the microorganisms could be introduced; (2) a ramp connecting the bridge piers and the deck. In addition, the upper part of the bridge was enclosed in a 5 mm wall to minimize lateral mycelial spread. The evaluation of this initial model showed the mycelium developing in all directions, indicating that the wall was not sufficient to contain growth (**Supplementary Figure 1B**). Following this initial test, several iterations based on observation-driven modifications were evaluated. This included comparing various surface topologies [smooth, diamond shaped (Gimeno et al., 2021), and undulating] that were implemented on the ramp and deck to assess their impact on fungal colonization. As surface topology did not impact colonization, the smooth design was selected to facilitate the printing process (**Supplementary Figure 1C1**).

To contain the mycelium within the wells, further improvements were implemented. For example, non-tilted printing resulted in a single smooth printed layer, ensuring the bridge’s contact surface with the multiplate. This feature was implemented to avoid a space that could be used by the mycelium for spreading beyond the wells. In addition, a continuous wall was implemented to enhance containment (**Supplementary Figure 1C2**). Despite these efforts, mycelia still grew beyond the two connected wells, spreading laterally. A final attempt (**Supplementary Figure 1C3**) focused on further restricting the mycelial spread by fitting the bridge directly to the multiplate cover. This involved reducing the lateral wall height and precision sculpting to ensure the cover closes properly. However, growth was still unconstrained as shown by the detection of microbial activity on adjacent wells (**Supplementary Figure 1D**). Therefore, the design approach was shifted to focus on a more constrained design, while still aiming to improve throughput.

Since well size was not an issue, we tested individual 2-well devices containing wells of the same dimension as in the 96-well plate that were connected by a bridge integrated directly within the device. In the first test, isolating the two wells was not sufficient to constrain mycelial development to the device; instead of using the crossing, the mycelium grew upwards reaching the Petri dish cover in which the device was placed (**Figure 1A1**). A second attempt with the same design but blocking upwards development by using hydrophobic PTFE tape wrapped over the inoculation well also failed, as the hyphae still grew on the surface of the device (**Figure 1A2**). Finally, by enlarging the crossing and making it closer to the inoculation point, we were able to constrain the growth within the device (**Figure 1A3**). With the concomitant use of the PTFE tape, the mycelia-forming organisms successfully used the crossing and reached the target well. To ensure safe and sterile use, an accompanying Petri dish-sized device holder was designed to accommodate ten 2-well units simultaneously (**Figure 1B**). The final device design allowed for running multiple parallel experiments with different organisms and/or media (**Figure 1C**). Considering the required manual handling, particularly when covering the device with PTFE tape of inoculation wells, we only detected one clear instance of human contamination (*Alcaligenes faecalis*; see later), suggesting that the devices can be handled without contamination. Overall, the individual 2-well device offered a good compromise between experimental handling and the increase in throughput that motivated the development of a new device.

### Validation of the method with model FH pairs in high and low nutrient conditions

To validate that diverse FH interactions could be studied with the devices, we started by testing 20 bacterial-mycelial organism pairs selected based on previous knowledge concerning their capacity to form FH (see **Table 1** and **2**). The experiments were performed using MA or MB in the inoculation and target compartments, respectively. Two Ascomycota (*T. rossicum* and *F. graminearum*), a Basidiomycota (*C. cinerea*), and the fungal-like oomycete (*P. ultimum*) were compared as FH-forming organisms. Those were paired with four bacterial species. Unfortunately, one of the fungi (*T. rossicum*) had to be excluded from the analysis because it sporulated profusely, which led to cross-contamination between individual devices. At the end, 15 combinations were compared. For all the FH-forming organisms, triplicate blanks without bacteria were also prepared. The sampling for culturing to validate bacterial growth was performed after confirmation by stereoscopy that the mycelium had crossed via the canal to the target well. In MA/MB media, this took place 15 days post-inoculation. None of the blanks yielded bacterial growth in the inoculation or target wells (**Figure 2A**).

**Figure 2.**
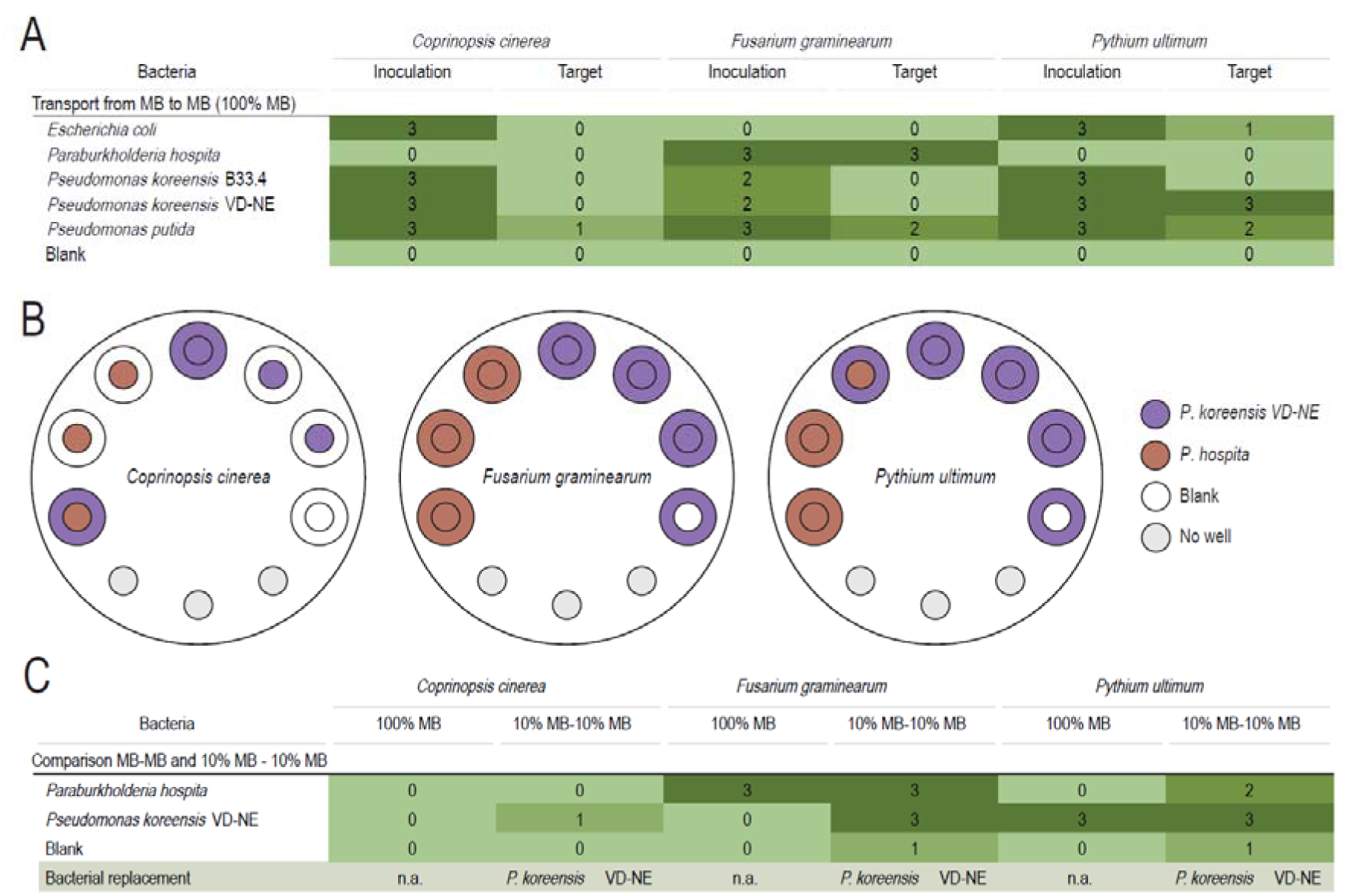
Testing of the devices using two media types. Shades of green indicate consistent crossing (darker) to no crossing (lighter) across three independent device replicates. **(A)** Rich medium (100% MB): Three mycelia-forming species and five associated bacterial partners were evaluated. Results are shown after two weeks of incubation. "Inoculation" refers to bacterial persistence in the inoculation well, while "Target" indicates bacteria detected in the target well. The “Blank” line in the table corresponds to our control wells in which only the fungus was inoculated and thus where no bacteria were expected. **(B)** Layout of the wells and bacterial replacement in the target wells for 10% MB. The smaller circle indicates the expected outcome in the target medium, corresponding to the inoculated bacterial strains. The bigger (outer) circles indicate the actual outcome. Same colored circles indicate that expected and actual outcomes were the same. Empty outer circles indicate no bacterial crossing. **(C)** Comparison of transport between rich (MB-MB) and poor medium (10% MB-10%MB). Only data from the target well is presented. Two bacterial partners (out of five tested) were assessed in combination with the same FH-forming species. Results correspond to sampling and bacterial culturing after three weeks of incubation. The “Blank” line in the table corresponds to control wells, in which only the fungus was inoculated and thus where no bacteria were expected. The “bacterial replacement” line indicates which bacterial strain replaced the inoculated bacteria; n.a.= not applicable.

The survival of the bacterium in the inoculation well was evaluated by re-culturing. For *C. cinerea* and *P. ultimum*, the bacterium *P. hospita* could not be recovered, while the other bacterial strains survived in all the replicates. For *F. graminearum*, only *P. hospita* and *P. putida* were consistently recovered (all three replicates) in the inoculation well. The other two *Pseudomonas* strains (*P. koreensis* VD-NE and *P. koreensis* B33.4) survived in two out of the three replicates. The bacterium *E. coli* was not recovered in any of the replicates in association with *F. graminearum* (**Figure 2A**). Next, we determined FH transport by culturing bacteria from the target well. The best FH transport rate was observed with the oomycete *P. ultimum* in which three bacterial strains (*P. koreensis* VD-NE, *P. putida*, and *E. coli*) were found in at least one replicate. The highest efficacy was observed with *P. koreensis* VD-NE (all replicates), followed by *P. putida* (2 of 3 replicates). Finally, *E. coli* was detected in one of the three replicates. Neither of the other two strains (*P. koreensis* B33.4 or *P. hospita*) were transported on FH of *P. ultimum*. Likewise, these two bacterial strains were not transported by *C. cinerea*. The only instance of FH with the fungus *C. cinerea* was for the transport of *P. putida* in one replicate. Bacterial colonization in the target well was not observed for any of the other bacterial strains with *C. cinerea* as FH. For *F. graminearum*, two of the bacterial strains tested were transported: *P. hospita* (all replicates) and *P. putida* (2 of 3 replicates) **(Figure 2A).** Together, the results demonstrated that the devices are useful to show FH associations, but transport in high nutrient media was overall poor.

The same pairs were then inoculated in diluted medium (10% MB) to compare transport under low nutrient conditions. This allowed us to investigate the impact of the nutrient conditions on the establishment of FH and the transport of bacteria. As survival in the inoculation well was not an issue, only the detection of bacteria in the target well is presented for clarity. Once more, the fungus *T. rossicum* had to be excluded from the analysis because of excessive sporulation, which led to *P. koreensis* VD-NE replacing all the other bacteria in the target wells (**Supplementary Figure 2**). Only the data for *P. hospita* and *P. koreensis* VD-NE are shown, because for the other combinations, mycelial growth was slower and the medium in the inoculation wells dried before the mycelium had crossed to the target medium. Indeed, in 10% MB, the incubation had to be extended to three weeks as crossing of the mycelium was not observed before this time. In this experiment, one blank without bacteria was included.

The comparison of FH transport under high and low nutrient conditions showed that the outcome was pair specific. For *C. cinerea*, which did not transport *P. hospita* or *P. koreensis* VD-NE in MB, low nutrient conditions stimulated FH transport of *P. koreensis* VD-NE in one replicate, and the same bacterium was detected as replacing *P. hospita* in another replicate (**Figure 2B** and **C**). For *F. graminearum* strain M47, FH transport of *P. hospita* was as efficient in 10% MB as in 100% MB. At 10% MB, the FH transport of *P. koreensis* VD-NE by *F. graminearum* was stimulated, and the bacterium was also detected in the blank (**Figure 2B** and **C**). As for MB, *P. ultimum* was the best FH transporter in 10% MB, with *P. hospita* crossing twice and *P. koreensis* VD-NE crossing multiple times. In one of the replicates, *P. koreensis* VD-NE replaced *P. hospita*, and in addition, it was detected in the blank (**Figure 2B** and **C**). Overall, the results demonstrated that the devices can be used to investigate FH between different nutrient conditions. Moreover, the results with the poor medium suggest a stimulation of FH, despite slower fungal growth. In addition, the detection of *P. koreensis* VD-NE replacing another bacterium or in the blanks suggests an increase in the exploratory behaviour of the mycelium under poorer nutrient conditions, which we explicitly investigated in the next series of experiments.

### Evaluation of increased exploratory growth under low nutrient conditions

To evaluate if the decrease in nutrient concentrations results in an increase in the exploratory growth of the fungus and facilitate the transport and replacement of bacteria to adjacent devices, we performed an experiment in which additional controls were included. For these experiments, *F. graminearum* M47 and *P. koreensis* VD-NE were selected. The experiments were performed in MA and 10% MA. The fungus or bacteria were placed individually in the inoculation well using a different layout (**Figure 3A**). The bacterial inoculum was placed either next to the fungus or surrounded by blanks with medium only. The devices were imaged after 4 dpi, but sampling was only performed after 7 and 11 dpi to avoid cross-contamination during sampling (**Figure 3B**). After 4 dpi, the fungus reached the target well in MA but had already traveled to the neighboring device in 10% MA. At 7 dpi, the fungus had colonized the neighboring devices in both nutritive conditions and had transported the bacteria to the target well in one of three replicates for 10% MA and two of three replicates for MA (**Figure 3C**). The results also showed that bacterial transport occur in poor and rich media. The same result was obtained after 11 dpi. In contrast, the bacteria are only detected in the inoculation well when inoculated in the vicinity to blanks. These results confirmed the hypothesis of an exploratory behavior allowing the colonization of the neighboring device, but the results do not support this to be more likely for low nutrient conditions (**Figure 3D**).

**Figure 3.**
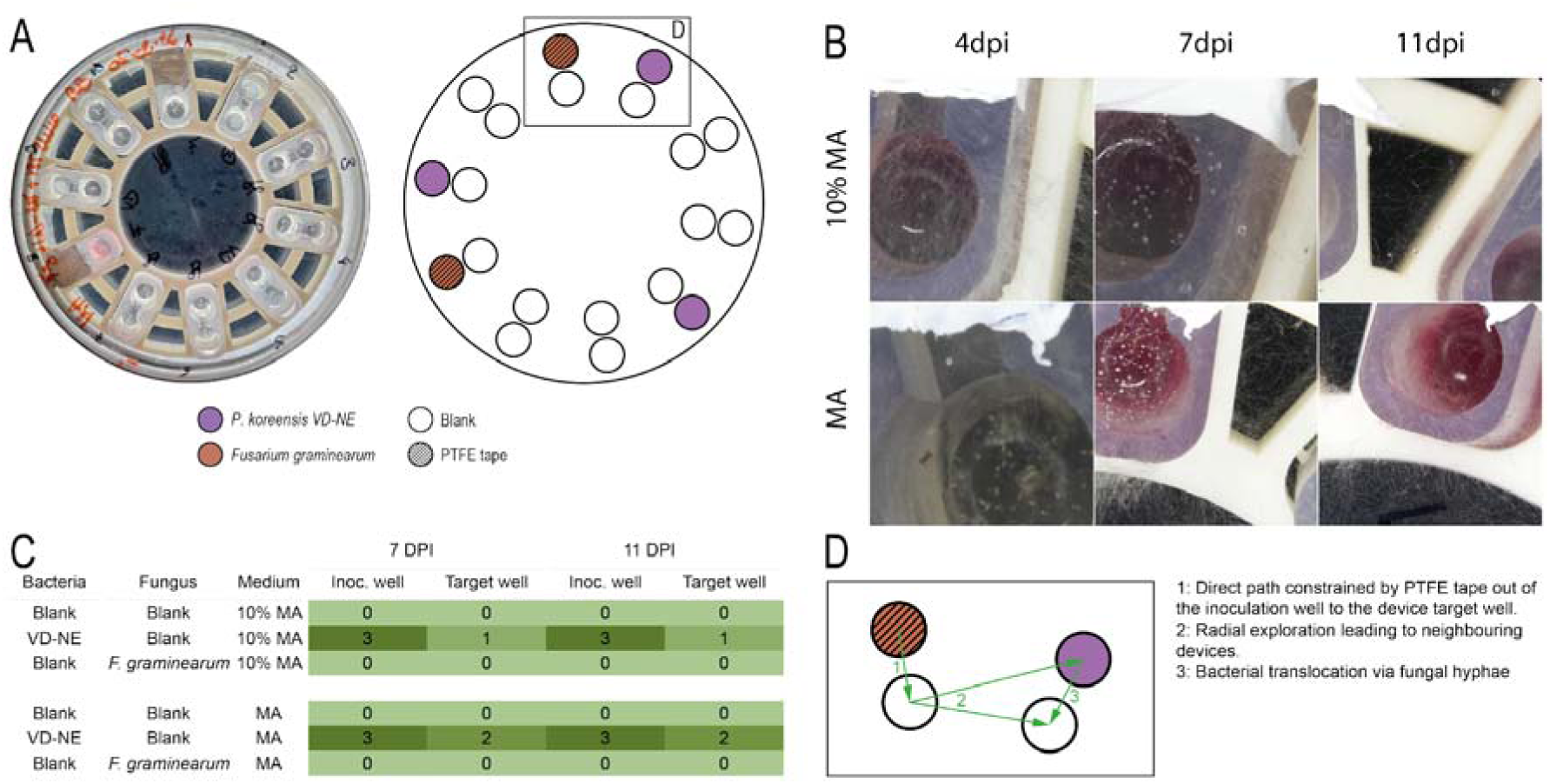
Evaluation of potential for bacterial replacement. For these experiments, *F. graminearum* strain K47 was used in combination with *P. koreensis* VD-NE as this couple was the most prominent in bacterial replacement events. The source well and target well contained MA or 10% MA. **(A)** Layout of the wells at the beginning of the experiment. In the scheme to the right, the outer circles indicate the inoculum while the inner circles corresponded to medium only at the start of the experiment. For this experiment, only two wells with fungi were considered to increase the number of blanks. **(B)** Pictures of the wells taken after 4-, 7- and 11-days post inoculation (dpi) in either MA or 10% MA. The pictures show that in less than 4 days in 10% MA and 7 days in MA, the fungus had crossed to the adjacent well and started exploring the neighboring devices. **(C)** Detection of the bacteria in the inoculation and target wells were recorded 7 and 11 dpi. Shades of green are used as in Figure 2. Experiments performed in triplicates. **(D)** Scheme of the putative path taken by the fungus, resulting in presence of bacteria in the target well of the device inoculated with bacteria only.

### Evaluation of the effect of the placement of the bacterial inoculum on FH

To further explore the effect of nutrients on FH, specific FH pairs studied with 10% MB were selected to test the impact of increasing heterogenous conditions (i.e., wells containing different nutrient conditions). For the tests with bacterial transport from poor to rich medium, we decided to again use *C. cinerea*, a "bad transporter" in the previous experiments and *F. graminearum*, a relatively fast-growing transporter. We coupled *P. hospita* to *F. graminearum* considering that consistent transport was obtained in rich medium (MB) while no transport was observed in poor medium (10%MB). In addition, *P. putida* was also included, as this bacterium was transported by all the FH-forming organisms in 100% MB. First, the pairs were inoculated together in the well containing 10% MB (cis-inoculation). The target medium corresponded to 100% MB. In the second experiment, the bacteria were inoculated in the 10% MB well, while the fungi were inoculated in 100% MB in the target well (trans-inoculation). The second condition mimicked the case in which the mycelia-forming species finds bacteria during exploration. Because drying of the wells was an issue for long incubations (e.g., in the nutrient-poor setting), in addition to the agar plug with the fungus, liquid medium (50 μl) was added in the wells to ensure higher humidity. The wells containing bacteria were filled, as described for the other assays, with 100 µl of the corresponding medium. Four replicates without bacteria were included in the assays, two of each corresponding to the cis or trans-inoculation in the case of *F. graminearum*.

In the experiment involving co-inoculation of *P. hospita* and *F. graminearum* in the same well (cis-inoculation) the bacteria survived in the inoculation wells until the last sampling points (13 days post inoculation, dpi). Bacteria crossed to the target well were detected as early as 4 dpi in two of three replicates. This number remained stable until 2 weeks post-inoculation (**Figure 4A** and **B**). Regarding the combination *P. putida*-*C. cinerea*, the bacterium was recovered in one of the replicates in the target well after 4 dpi. After 7 dpi, the bacterium was detected in an additional target well replicate. In the combination of *P. koreensis* with *C. cinerea* the bacterium did not cross for the duration of the experiment (**Figure 4A**).

**Figure 4.**
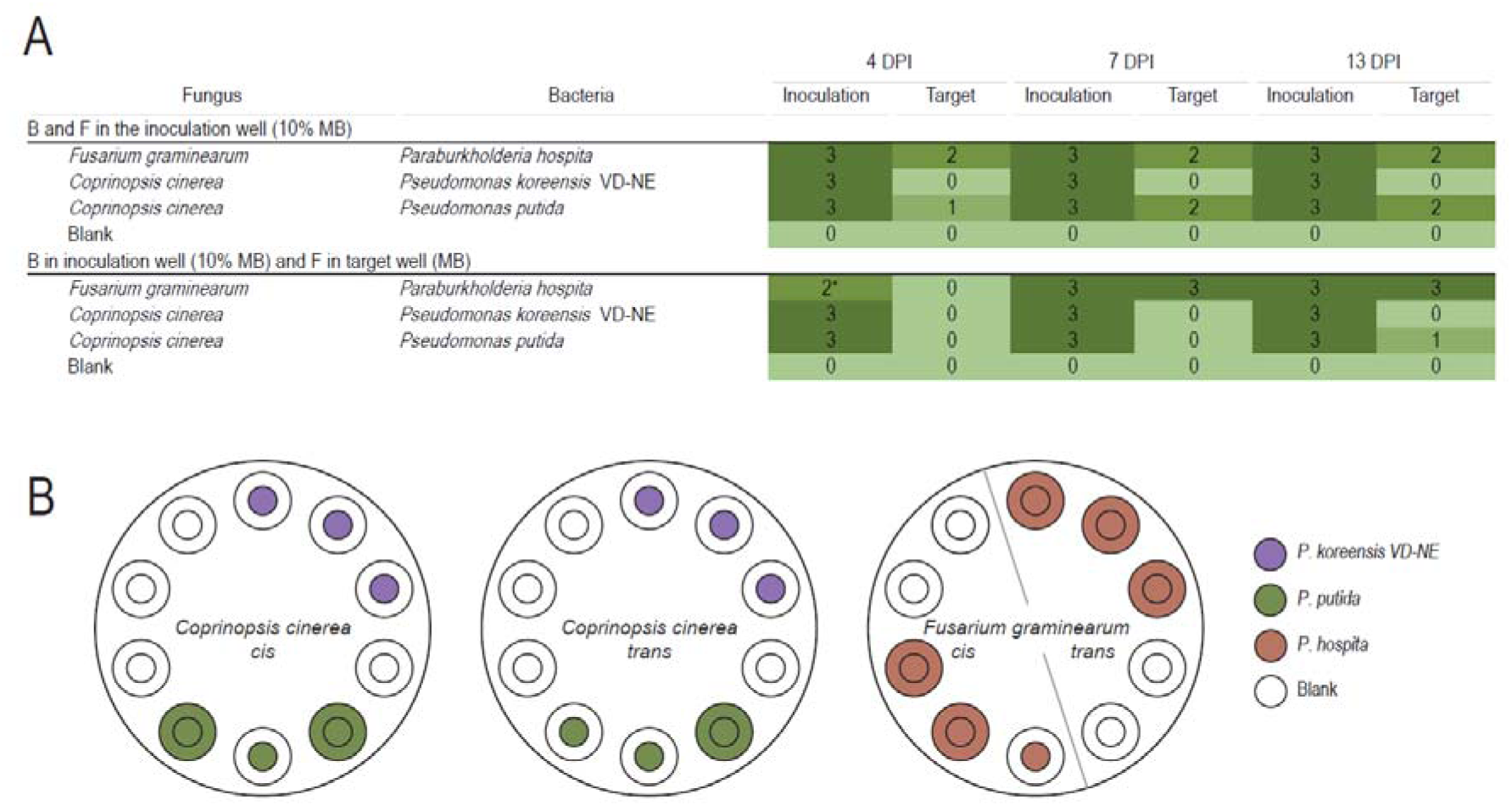
Assessing FH interactions with two inoculation modes and different nutrients. Shades of green are used as in Figure 2. Devices in triplicates**. (A)** Upper part: Co-inoculation in poor medium (10% MB) (cis-inoculation); two fungal species (*F. graminearum* and *C. cinerea*) were co-inoculated with the bacterial partners in the inoculation well, while the target well contained MB. Bacterial crossing to the target well was tested at 4, 7, and 13 dpi. Lower part: Inoculation in separate wells (trans-inoculation); bacteria were inoculated in the poor medium (10% MB) well, while fungi were placed in the adjacent rich medium (MB) target well. The “Blank” line in the table corresponds to control wells in which only the fungus was inoculated. The asterisk indicates a likely error in sampling. No bacterial crossing was observed at this time point. **(B)** Layout of the wells in the target wells after 13 days. The smaller circle indicates the expected outcome in the target medium (i.e., inoculated bacterial strain). The bigger (outer) circles indicate the actual outcome. Same colored circles indicate that expected and actual outcomes were the same. Empty outer circles indicate no bacterial crossing.

In the assay where the bacterium and the fungus were inoculated in opposite wells (the bacteria in the inoculation well with 10% MB and the fungi in the target well with MB), all the bacteria survived in the inoculation well except for one replicate of *P. hospita* (associated to *F. graminearum*) at 4 dpi. Since the same replicate showed survival later, we concluded that sampling at 4 dpi was potentially inadequate leading to a false negative. Despite fungal hyphae reaching the target well by 4 dpi, none of the bacteria were recovered in the target well at this sampling point. However, after 7 and 13 dpi, *P. hospita* had crossed on *F. graminearum* in all the replicates (**Figure 4B** and **C**). In contrast, in the trans-inoculation with *C. cinerea*, bacteria were only detected in one replicate (*P. putida*) after 13 dpi (**Figure 4B** and **C**). In addition, the bacterium was also detected in one of the blanks at 20 dpi (**Supplementary Figure 3**), but given the extended incubation period, this result might not bear any biological meaning and was not considered indicative of transport or replacement.

Overall, the results show that both configurations (trans- and cis-inoculation) promote the formation of FH. However, as expected, in the cis-inoculation, colonization of the target medium by the bacterium takes longer than when the bacteria and the fungi are co-colonized in the same initial medium. This also shows that the movement of bacteria does not necessarily follow the direction of fungal growth.

### Evaluation of the devices using non-model FH pairs

For the final experiments, we expanded the set of organisms to include bacteria and fungi that had not been used in previous FH experiments. We wanted to test the potential for FH associations between bacteria and fungi involved in necromass decomposition. First, we used strains belonging to genera reported in the necrobiome (Perez-Pazos et al. 2024; Cantoran et al. 2023) but not directly isolated from necromass. The conditions that showed a more promiscuous bacterial crossing with the model strains (10% MB in both inoculation and target wells and co-inoculation) were used for this experiment. One blank without bacteria was used per device. The bacterium *P. fluorescens* was consistently transported by *Paecilomyces* sp., *Mortierella* sp., and *T. atroviridae*. The bacterium *S. marcescens* crossed with the same fungi, albeit with different efficacy. Conversely, none of the bacteria crossed on FH produced by *C. globosum* (**Figure 5**). The bacterium *S. marcescens* was detected in the blank of *Paecilomyces* sp. Also, in one of the replicates with *T. atroviridae*, *Chryseobacterium* replaced *S. marcescens*. Overall, these experiments with non-model species showed that different fungal-bacterial combinations differ in their capability to form FH but also that even for species not previously known to form FH, this type of association could be demonstrated experimentally with the devices.

**Figure 5.**
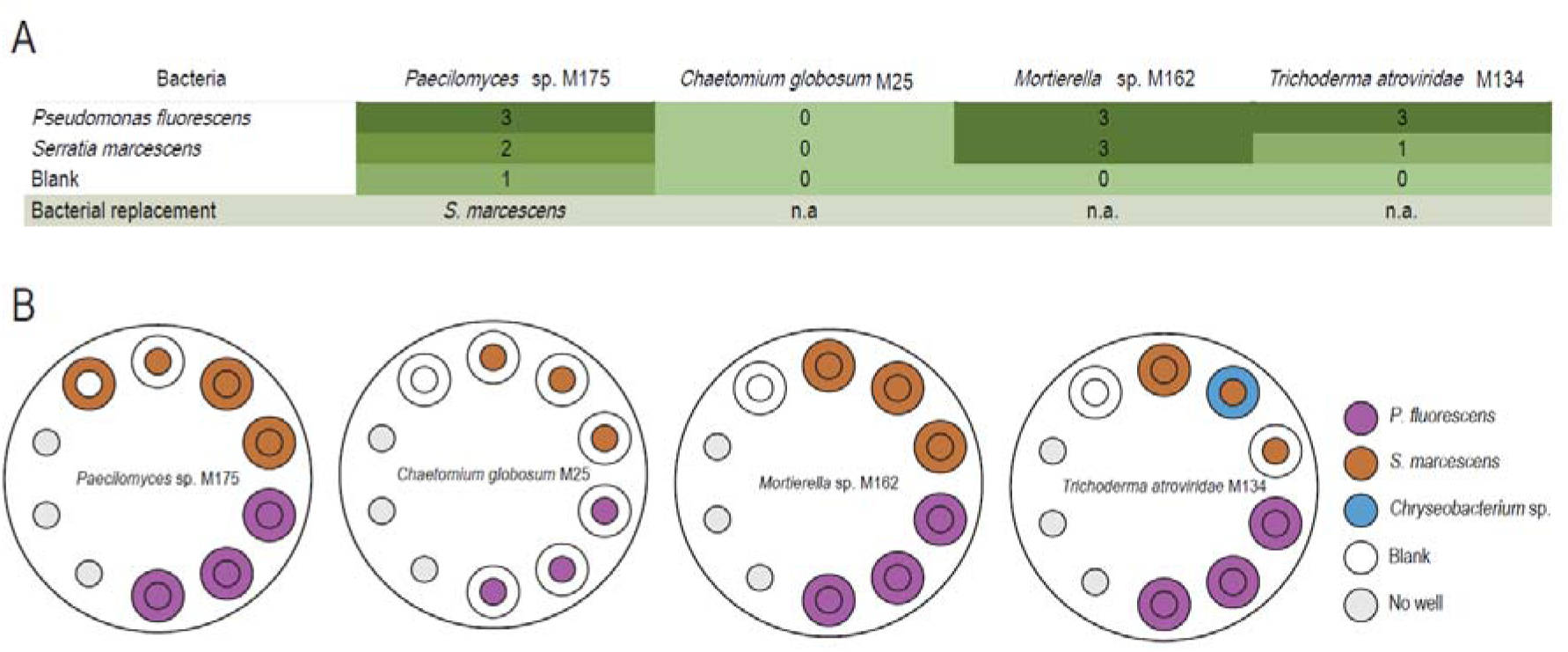
Evaluation of FH transport in nutrient-poor conditions using non-model strains. **(A)** Four fungal species were tested in combination with two bacterial partners in a homogeneous poor medium (10% MB in both inoculation and target wells). Experiments were conducted in triplicate, and the results of bacterial transport were recorded two weeks post-inoculation. The “Blank” line in the table corresponds to control wells, in which only the fungus was inoculated and thus where no bacteria were expected. The “bacterial replacement” line indicates which strain replaced the expected bacteria; n.a.= not applicable. Shades of green are used as in Figure 2**. (B)** Layout of the wells to illustrate the bacterial replacement in the target wells. The smaller (inner) circles indicate the expected outcome in the target medium (i.e., inoculated bacterial strain). The bigger (outer) circles indicate the actual outcome. Same colored circles indicate that expected and actual outcomes were the same. Empty outer circles indicate no bacterial crossing.

Finally, an experiment was performed using a more complex substrate (necromass with low and high melanin content) and strains isolated directly from necromass. The three conditions compared were 10% PDB and necromass with low (LMN) and high (HMN) melanin content as the medium for growth in the target well. The inoculation well always contained fungal plugs from a PDA plate with bacteria inoculated on top. A blank with the fungus alone was included.

In LMN, crossing in association with *Chaetomium* sp. 31 was observed in all replicates of *Flavobacterium* sp. and 1 out of 3 replicates of *Pseudomonas* sp.. The *Chryseobacterium* sp. and *Pseudomonas* sp. were recovered in the target well with 10% PDB in one of the replicates in association with *Chaetomium* sp. 31. *F. cupreum* did not cross in 10% PDB with this fungus. The blank of *Chaetomium* sp. 31 in LMN corresponded to the only instance of external contamination by detection of *A. faecalis*. No bacterial crossing was observed in HMN in association with *Chaetomium* sp. 31.

In the case of the second fungus, *Trichoderma* sp. 48, all the bacterial strains tested crossed in LMN. Moreover, the blank was replaced by one of the inoculated bacteria (*Pseudomonas* sp.). Only one of the replicates of *Chryseobacterium* sp. crossed in PDB 10% with *Trichoderma* sp. 48. In HMN, only one occurrence of transportation was found (*F. cupreum*) (**Figure 6**). The results with more complex substrates highlight the importance of the interplay between biological and biochemical factors on the establishment of FH and provides additional support for the importance of this type of interaction on the co-colonization of complex substrates commonly found in natural environments such as soils (Maillard et al. 2023).

**Figure 6.**
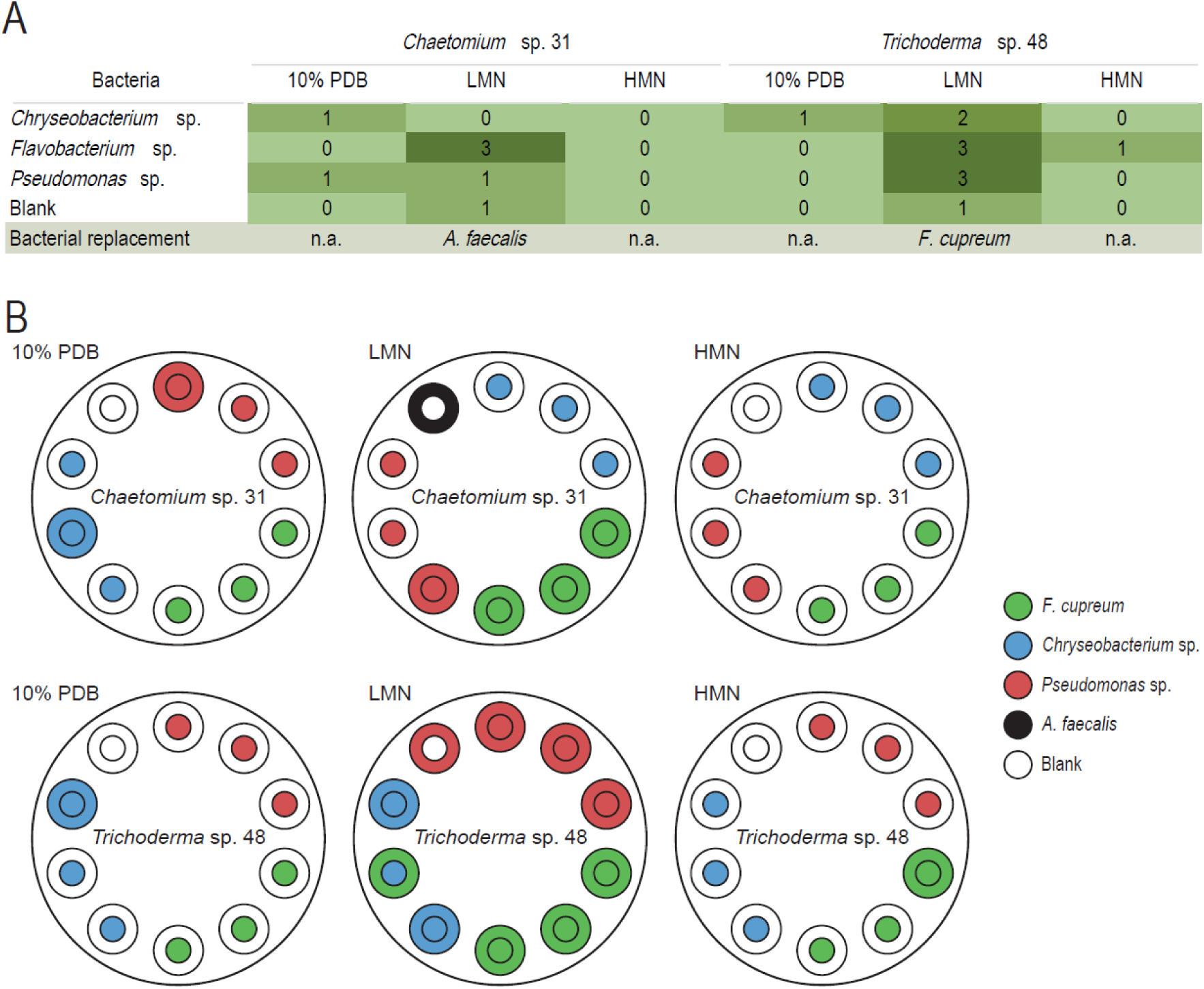
Evaluation of FH transport using necromass-based media. **(A)** Shades of green are used as in Figure 2. Two fungal species were tested in combination with three bacterial partners using three media types in the target well: 10% PDB, and two necromass-based media: LMN) and HMN. The source well and target well contained PDA. Experiments were performed in triplicates, and the results were recorded two weeks post-inoculation. The “Blank” line in the table corresponds to devices in which only the fungus was inoculated and thus where no bacteria were expected. The “bacterial replacement” line indicates which bacterial strain replaced the expected bacteria; n.a.= not applicable. **(B)** Layout of the wells and bacterial replacement in the target wells. The smaller circle indicates the expected outcome in the target medium, corresponding to the inoculated bacterial strains. The bigger (outer) circles indicate the actual outcome. Same colored circles indicate that expected and actual outcomes were the same. The dark outer circle indicates the only instance of human contamination.

### Comparison of the observations made in the 2-well devices with *in planta* transport of bacteria via FH

To test whether *F. graminearum* can transport *P. hospita in planta*, an approach was developed to assess fungal-mediated transport on and between wheat spikes. In field conditions, and particularly under saturated humid conditions caused by rain showers occurring before dawn, *F. graminearum* hyphae can grow on the wheat infected surface allowing the fungus to colonize neighboring areas of the spikelets (**Figure 7A**). This growth may enable the fungus to transport bacteria to neighbouring spikelets on the same spike, as well as to spikelets on adjacent wheat spikes, facilitated by wind-induced contact between spikes. An initial experiment was performed to determine whether the fungus can generate mycelia on the surface of wheat infected spikelets under controlled growth-chamber conditions. Two spikelets from a spike were inoculated with *F. graminearum* spores of the strain PH-1. Inoculated spikes were detached and placed inside a box under saturated humid conditions (**Supplementary Figure 4A**). Fungal growth outside the inoculated spikelet was observed 2 dpi, and by 5 dpi fungal hyphae fully covered adjacent spikelets (**Supplementary Figure 5**). Therefore, 5 dpi was selected for hyphae collection in subsequent co-inoculation experiments with bacteria. Next, a second experiment was done in which the fungus was simultaneously co-inoculated either with *E. coli*-RFP or *P. hospita*-RFP bacterial strains. *Escherichia coli* was selected as the negative control as the earlier experiments had shown that *E. coli* cannot disperse with *F. graminearum* in the 3D-printed device (**Figure 2A**). An inhibitory effect was observed when *F. graminearum* was co-inoculated with *P. hospita*-RFP (**Supplementary Figure 4B**). To test whether the fungus could transport bacteria, a third experiment was performed. Two spikelets from a wheat spike at the anthesis stage were inoculated with the fungus one day prior to bacterial inoculation to avoid the inhibitory effect from *P. hospita*-RFP (**Figure 7B**). Spikes containing the two spikelets inoculated first with the fungus and then with the bacteria were detached from the plant and incubated in a humid chamber. To increase the amount of tissue colonized by hyphae and therefore increase the likelihood of bacterial transport and detection, an uninfected wheat spike was positioned in front of the inoculated spikes (**Figure 7B and 7C**). To track the bacterial strain beyond the two inoculated spikelets, *E. coli*-RFP and *P. hospita*-RFP carrying a plasmid expressing RFP and conferring resistant to gentamicin were used. Resistance to gentamicin allowed the selection of bacteria on agar plates and RFP accumulation produces a reddish coloration to distinguish the test bacteria from other bacterial species or strains. At 5 dpi, infected tissue and hyphae growing beyond the two co-inoculated spikelets were collected, macerated and an aliquot of the suspension was plated onto agar plates containing gentamicin and cycloheximide. The two co-inoculated spikelets were separately collected and macerated to confirm the presence of bacteria during the whole experiment. As a control, the fungus was co-inoculated with water (**Supplementary Figure 6**). Bacteria were detected in 1 out of 3 replicates when the fungus was inoculated with water. However, the isolated colonies lacked the characteristic reddish color of either *P. hospita*-RFP or *E. coli*-RFP, and PCR amplification of the *RFP* gene was negative (**Figure 5E and Supplementary Figure 6**). In contrast, in the co-inoculated spikelets *E. coli*-RFP was detected in 2 out of 3 replicates, and *P. hospita*-RFP was detected in all three replicates and the colonies recovered from those plates exhibit the reddish coloration and were positive for RFP amplification (**Figure 7D and 7E, and Supplementary Figure 6**).

**Figure 7.**
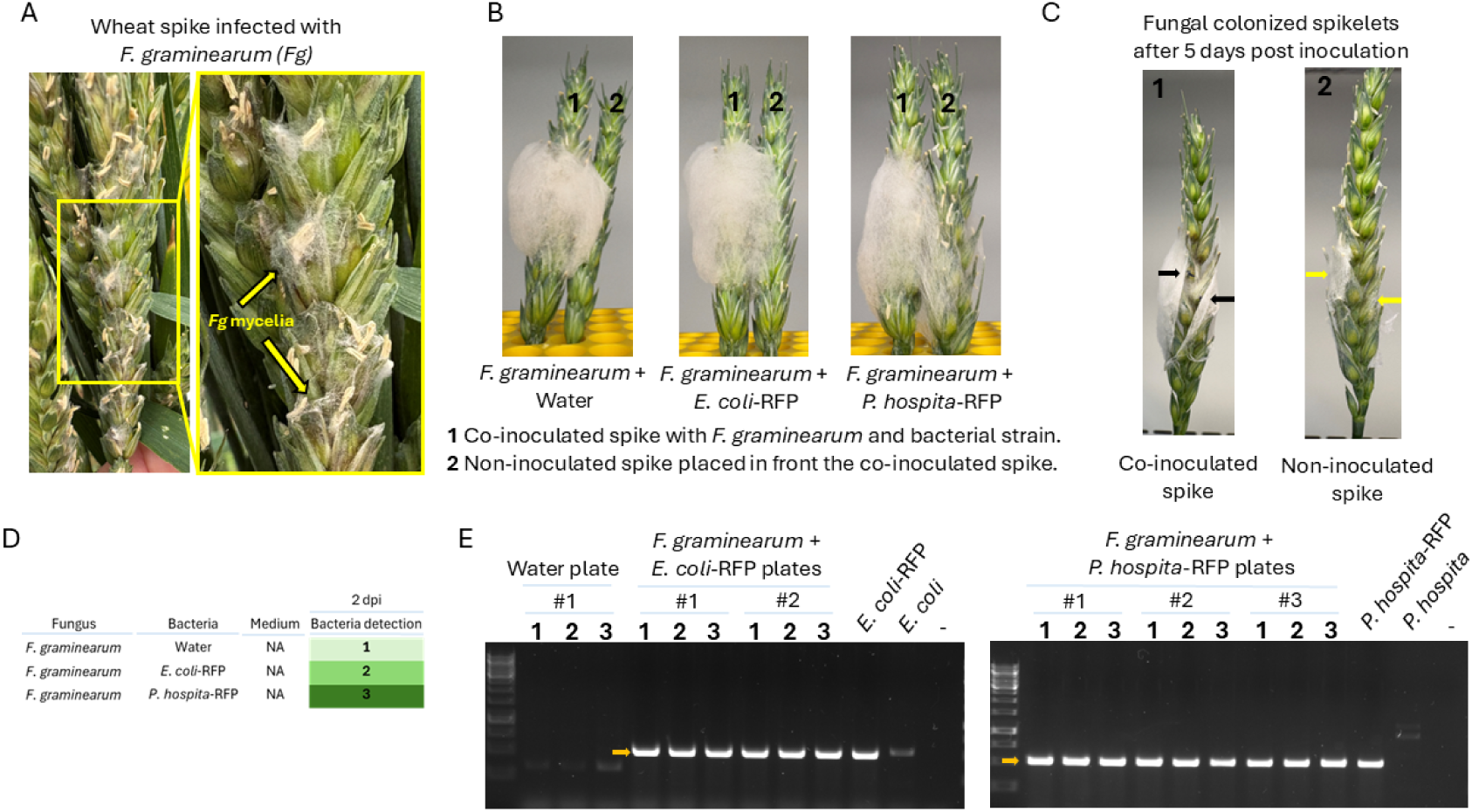
Evaluating *in planta* bacterial transport by *F. graminearum*. (**A**) Wheat spikes inoculated with *F. graminearum* UK-strain Fg602 under field conditions (left panel). Right panel is a magnification from the left panel showing mycelia growing around the anthers after a short period of rainfall and cloudy weather (yellow arrows). (**B**) Wheat spikes co-inoculated with *F. graminearum* strain PH-1and bacterial strains under controlled growth-chamber conditions. Fungal inoculation prior to bacterial inoculation prevents the negative impact that *P. hospita* exerts on fungal growth. Images were taken 5 dpi. Number 1 denotes the spike where two spikelets were co-inoculated with the fungus and bacteria while number 2 denotes a non-inoculated spike that was placed in front the inoculated spike to promote fungal colonization and increase the likelihood of bacterial transport. (**C**) After 5 dpi, left panel: colonized spikelets below the point of co-inoculation (black arrows) as well as where mycelia were collected. In addition, the two co-inoculated spikelets were separately collected. Right panel: Mycelia and colonized spikelets (yellow arrows) from the spike placed in front the co-inoculated spike were also collected. (**D**) Number of replicates in which bacteria were detected beyond the co-inoculation site. Three technical replicates per combination were performed. (**E**) Diagnostic PCR for *RFP* gene amplification. Bacterial colonies recovered from macerates of mycelia and colonized spikelets beyond the co-inoculation site were evaluated by PCR. Bacteria recovered from a plate where the fungus was co-inoculated with water (Water plate) were negative for PCR amplification while bacterial isolates recovered from plates where the fungus was co-inoculated either with *E. coli*-RFP or *P. hospita*-RFP were positive. Numbers denote PCR evaluation of three out of six colonies isolated from each plate (replicate). “-” refers to a PCR without template. *E. coli* and *P. hospita* strain lacking the pBBR-RFP plasmid were included to test primer specificity. Expected amplicon size: 538 bp. A faint, slightly larger nonspecific PCR band was observed for the *E. coli* strain.

To confirm bacterial transport by *F. graminearum*, the experiment was repeated a second time. In addition, spikes inoculated only with *P. hospita*–RFP or *E. coli*–RFP were included to rule out cross-contamination during the inoculation and sampling procedures. Spikelets from the spike positioned in front the co-inoculated spike were collected and processed separately to determine the localization of the bacteria. Bacteria were not detected beyond the inoculation site in spikes inoculated solely with *P. hospita*–RFP or *E. coli*–RFP, indicating that these bacterial strains cannot independently colonize adjacent spikelets. Furthermore, no bacteria were detected in spikelets inoculated only with the fungus **(Figure 8A).** However, bacteria were detected beyond the inoculation site when co-inoculated with the fungus. *E. coli*–RFP was detected in 2 out of 3 replicates, and *P. hospita*–RFP was detected in all three replicates. Both bacterial strains were also detected in spikelets recovered from the adjacent non-inoculated spike in only one replicate, suggesting that their localization is mainly confined to spikelets recovered from the spike where co-inoculation with the fungus occurred **(Figure 8A).** Finally, bacteria recovered from those plates show the reddish coloration due to RFP expression and were positive for RFP amplification **(Figure 8B and Supplementary Figure 7).** Altogether, these results indicate that *F. graminearum* could transport both bacterial strains.

**Figure 8.**
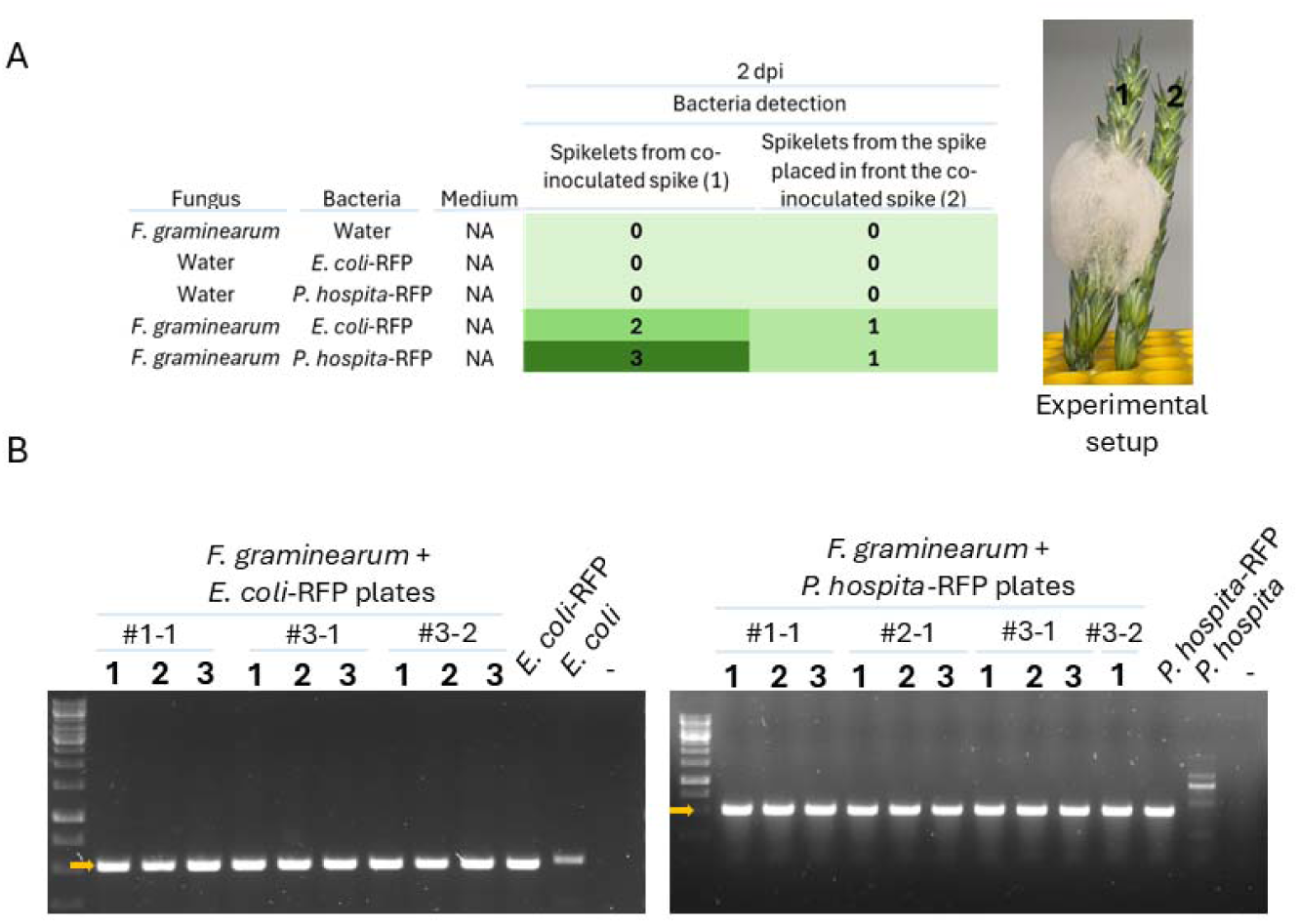
*F. graminearum* can transport bacteria *in planta*. (**A**) Number of replicates in which bacteria were detected. Tissue colonized by the fungus was divided into two groups: spikelets from the spike where the fungus and bacteria were co-inoculated, and spikelets from the non-inoculated spike placed near the inoculated wheat spike. Three technical replicates per combination were performed. (**B**) Diagnostic PCR was positive for bacterial isolates recovered from plates containing macerates of colonized tissue and mycelia of *F. graminearum* + *E. coli*-RFP and *F. graminearum* + *P. hospita*-RFP. #1-1 and #2-1 refer to independent replicates in which macerates of mycelia and colonized spikelets beyond the co-inoculation site were plated. Each replicate was divided in two as spikelets from the co-inoculated spikes (#3-1) and spikelets from the spikes placed in front the co-inoculated spike (#3-2) were processed separately. Numbers denote PCR evaluation on colonies isolated from each plate (replicate). “–” refers to a PCR reaction without a template. *E. coli* and *P. hospita* strains lacking the pBBR-RFP plasmid were included to test primer specificity. Expected amplicon size: 538 bp. A faint, slightly larger nonspecific PCR band was observed for the *E. coli* strain.

## Discussion

Dispersal, drift, selection and speciation are fundamental ecological processes driving the structure of a community by modulating its generation and maintenance. Despite the widely accepted “everything is everywhere” notion of microbial biogeography, mounting evidence suggests that single celled microbes are dispersal limited. While dispersal is a fundamental ecological process, the challenges with its quantitative assessment have affected the way its impact can be appreciated in microbial ecology (Custer et al., 2022). The aim of this study was to propose a novel way to evaluate dispersal of bacteria in the context of FH. The devices presented in this study offer new insights into this specific mechanism in which the dispersal of motile bacteria is promoted using the networks of mycelial organisms (Furuno et al., 2010; Kohlmeier et al., 2005). This mechanism of dispersal combines two active microbial dispersal processes: hyphal growth and flagellar motility but could potentially involve more passive processes due, for instance, to water redistribution associated with hyphal growth (Clark et al., 2024; Richter et al., 2022). The experimental tool developed here can complement existing systems to contextualize and quantify the importance of FH on the structuring and function of microbial communities under dispersal limitation.

To overcome the limitations of existing systems to evaluate FH in the laboratory, we developed a 3D-printed device designed to increase throughput while maintaining flexibility in medium composition. Unlike existing approaches, which often restrict the number of simultaneous assays or require specialized setups, this device enables parallel testing of up to 10 FH pairs under varied nutritional conditions. Its modular design facilitates easy customization and reproducibility, while the use of 3D printing ensures cost-effectiveness and rapid prototyping. The devices were used here to assess colonization. Nonetheless, by isolating factors such as nutrient bioavailability in the inoculation and target medium, timing of partners meeting and partner traits, we can gain a richer understanding of how substrate heterogeneity and priority effects shape community assembly (Fukami, 2015). The use of the devices offers insight into the effect of trophic factors and the physical environments on FH dispersal into the target medium as a first step to disentangle the processes that lead to the successful establishment of microbial communities in a given habitat (Custer et al., 2022).

### Validation of the devices and sources of variability

Our findings demonstrate the potential of the 3D-printed device for screening mycelial–bacterial associations in the context of FH. The design provides a controlled setting for dispersal studies. Despite multiple cycles of design and improvement, variability across replicates persisted, indicating that additional factors beyond environmental control influence bacterial movement. Desiccation emerged as a likely contributor (Kuhn et al., 2022), as water availability is critical for forming thin films that enable bacterial movement along hyphae (Aung et al., 2018; Dechesne et al., 2010). In the context of FH, the ability to modify traits such as motility, stress resistance, or interaction mechanisms will play a significant role on the successful colonization of a novel environmental niche (Buffi et al., 2025; Fischer & Glass, 2019; Jia et al., 2024; Jiang et al., 2021). These variations could explain why some replicates showed delayed or inconsistent colonization of the target well. Furthermore, intra-population differences in bacterial motility may explain inconsistent results, as a previous study reported substantial variation in dispersal speed within populations. Indeed, it has been shown that individuals within *P. putida* populations expressed substantial differences in motility and speed, with some cells being slow-moving or even static while others could reach high velocities (up to 100 µm/s) (Buffi et al., 2025). Those differences could represent bet-hedging strategies allowing the populations to efficiently colonize different types of environments. Moreover, our observations indicate that bacterial dispersal was not strictly directional; when bacteria reached the target well, they were almost always still present in the inoculation well, suggesting a pattern of expansion rather than relocation. These findings highlight the complexity of FH and suggest that extended incubation times or refined moisture control could improve reproducibility.

While our device offers intermediate throughput and flexibility, it requires attentive manipulation. Due to the amount of handling required, particularly during sampling and the application of PTFE tape, it could be more susceptible to contamination than simpler designs such as fungal drop assays (Buffi et al., 2023) or split agar systems (Junier et al., 2021). Only one clear instance of human-derived contamination was detected, suggesting that the use of careful sterile techniques can be sufficient to limit this issue. However, an alternative design for long incubations could be the use of destructive sampling considering one (or multiple) devices per time point. This is possible given the ease of printing and preparation of the devices. Another potential limitation of the design is that the presence or absence of bacteria is determined by sampling the wells and re-culturing on a generic medium (e.g., nutrient agar) supplemented with the antifungal cycloheximide. During sampling, insufficient homogenization may result in missed detection. Accordingly, sampling must be performed carefully to ensure reliable results, and we encountered one instance of sampling error in our assays, which led to a false negative for the detection of bacteria (detection of *P. hospita* at 4 dpi in the inoculation well in the experiment with *F. graminearum*). However, we argue that when sampling is done properly, specifically by thoroughly homogenizing the wells through repeated pipetting up and down, the results are consistent, as demonstrated by reproducible outcomes across multiple sampling days. Additionally, pipetting can damage hyphae, mobilize spores, and impose mechanical stress, potentially altering fungal–bacterial interactions. We also observed an increase in bacterial replacement events (see below) in the experiments involving several rounds of sampling. For these reasons, minimizing handling and avoiding highly sporulating fungi are recommended to improve reproducibility.

### Impact of medium and inoculation strategy

Our results underscore the critical role of medium composition and inoculation configuration in shaping bacterial movement along hyphae. For instance, *C. cinerea* generally performed poorly as a highway for *Pseudomonas* strains, yet the two-well device with varying nutrient conditions, two out of three replicates of *P. putida* KT2440 successfully crossed when co-inoculated with the same fungus. This highlights how both abiotic conditions and inoculation strategy influence FH outcomes. Differences in medium composition may also explain discrepancies with previous studies. Junier et al., (2021) reported FH associations between *Pseudomonas* spp. and *C. cinerea* in a heterogeneous matrix, contrasting with our findings in more uniform conditions. Similarly, Nazir et al., (2014) showed broad dispersal of *P. terrae* along fungal hyphae of *Lyophyllum* sp. strain Karsten and *Fusarium oxysporum* in soil-based media, whereas the closely related strain, *P. hospita* (Pratama et al., 2020), displayed variable success depending on medium richness and mycelial partner. Interestingly, in heterogeneous set-ups, *P. hospita* crossed more frequently when inoculated opposite the fungus (trans-inoculation) rather than co-inoculated (cis-inoculation), suggesting that fungal colonization of the target medium may enhance transport potential or reduce initial competition that could impair the establishment of FH.

We also observed a link between fungal exploratory behavior and bacterial translocation. An intriguing observation made with the devices was bacterial replacement (meaning a bacteria inoculated in a different device replacing the expected bacteria in the target well). This was especially common with fast-growing fungi like *F. graminearum* when inoculated in poor media. In contrast, inoculation in rich media reduced both bacterial replacement and crossing events. These contrasting results are likely driven by enhanced fungal exploratory behavior under nutrient limitation (Camenzind et al., 2022; Heaton et al., 2015), something that was particularly evident for *F. graminearum*, a highly exploratory fungus (Ejaz et al., 2023; Todorović et al., 2023). The experiment in which bacteria were placed in an adjacent device clearly shows that replacement is likely the result of lateral fungal exploration rather than contamination (**Figure 3**), something that should be considered during the experimental design.

The changes in fungal exploratory growth may reflect the impact of environmental cues and challenging conditions on fungal stress responses and hyphal expansion (Camenzind et al., 2022; Fukasawa & Ishii, 2023), something that bacteria might exploit. Flagellated bacteria, for example, may use fungal stress signals such as ethylene, known for inter-kingdom communication (Bidon et al., 2020; Papon & Binder, 2019; Shekhawat et al., 2023), to modify the direction of hyphal growth and favor dispersal when resources are scarce. Flagellated movement is active and thus costly (Kuhn et al., 2022; Pion et al., 2013) and might be regulated by available resources. Phenotypic variation within bacterial populations could further influence dispersal, with nutrient-deprived phenotypes responding faster to chemotactic signals. The impact of fungal exudates and volatile organic compounds, previously identified as key drivers of bacterial–fungal interactions (BFI) (Deveau et al., 2018; Palmieri et al., 2019), warrant deeper investigation to clarify their role in the type of FH dynamics observed here (Deveau et al., 2018; Shamugam & Kertesz, 2023).

To better characterize bacterial replacement events, we separately inoculated *P. koreensis* VD-NE and *F. graminearum* strain M47, ensuring that bacterial crossing could only happen from the growth of the hyphae to a bacterial well and a further crossing to a different well.

Bacterial replacement happened twice in MB and once in 10% MB after 7 days (**Figure 3**). These observations imply that the fungus could reach other wells and allow the displacement of bacteria relatively far from its inoculation point. These results confirm that bacterial replacement likely happened in other experiments considering the potential for the fungal partner to cross to neighbouring wells and transport bacteria. In this assay, no significant difference in crossing was observed between the two media. A fundamental factor modulating medium influence could be spatial arrangement of the partners. Our observations on bacterial replacement highlight the need for tagged bacteria or identification of colonies with DNA barcoding as microorganisms could move further than their designated target well. This instance could prove very useful in further study as it demonstrates transport to different distances, potentially modulated by spatial arrangement, nutrient content, and timing of partners meeting.

These results also emphasize the need to distinguish between transport upon co-inoculation or replacement favored by hyphal exploration. Bacterial replacement, for instance, could reflect dispersal in response to specific environmental cues. For example, (Otto et al., 2017) demonstrated that *Bdellovibrio bacteriovorus* disperses along fungal hyphae only when prey is present, highlighting cue-dependent movement behavior. Our findings therefore call for further investigation into the ecological and functional consequences of selection and replacement during exploration, as well as the mechanisms regulating these processes. We believe that our devices provide a promising framework to explore the drivers of active bacterial dispersal, as they enable controlled modulation of key factors, including nutrient availability.

### Influence of necromass as a resource in FH

Another factor that can mediate FH is substrate quality. We used fungal necromass to explore how resource bioavailability influences bacterial movement along fungal hyphae (Beidler et al., 2020, 2025; Fernandez & Kennedy, 2018; Maillard et al., 2023). Our results reflected known constraints associated with necromass degradation: five out of six bacterial–fungal pairs crossed in LMN, whereas none crossed in HMN. LMN appears to support higher bacterial activity based on 16S rRNA gene copy counts (Maillard et al., 2025), providing sufficient energy for active movement along hyphae, while HMN limits nutrient accessibility. Similarly, fungi grow faster and produce more biomass on LMN given its higher quality as a nutritional resource (Beidler et al., 2020; Fernandez & Kennedy, 2018, Maillard et al., 2025) and thus are more likely to support a bacterial partner. Furthermore, melanin content seems to more strongly affect fungal growth than bacterial growth in liquid media (Narayanan et al., 2025; Haq et al., 2024; Novak et al., 2024) suggesting that fungal growth may govern the strength of the association. Hydrophobicity of the substrate may also play a role: fungal melanin is hydrophobic (Suthar et al., 2023), and fungi can modulate cell surface hydrophobicity via molecules such as hydrophobins (Bayry et al., 2012). The dynamic interactions between the substrate and the hyphal surface may influence attachment and movement, as closer contact between hydrophobic surfaces reduces the energetic cost of water exposure (Sun, 2022). Together, these findings highlight the importance of resource bioavailability and surface properties in shaping FH dynamics.

### Bacterial transport during *F. graminearum* colonization of a cereal host niche

*F. graminearum* is a major causal agent of Fusarium Head Blight (FHB) on wheat, barley, and rice (Wegulo et al., 2015). Wheat spikes are most susceptible to FHB at anthesis, when the florets inside the spikelets are open, allowing the fungus to penetrate host tissues and colonize the entire spike. Under warm and humid conditions, the fungus can develop pinkish mycelia on the surface of infected tissues (Alisaac & Mahlein, 2023) (Figure 7A), which can potentially be subsequently used by bacteria as FH. To exploit this natural host colonization biology and thus study bacterial transport by *F. graminearum*, we developed an approach focused on wheat spike colonization. We observed that *F. graminearum* generates abundant mycelia on inoculated wheat spikelets under saturated humidity conditions. The mycelia grow from the inoculated spikelets toward non-linoculated ones, both within and on the surface of the host tissue, which proved useful for testing bacterial transport beyond the inoculation site. Additionally, inoculating the bacteria directly inside the wheat floret reduces the chance of bacterial cross-contamination. When bacterial strains were inoculated alone, bacteria were not detected beyond the inoculation site, confirming that this approach is suitable for studying fungal-mediated bacterial transport under *in planta* conditions.

A higher number of bacteria beyond the inoculation site was detected in the first experiment compared with the second experiment for both *E. coli*-RFP and *P. hospita*-RFP. This difference may be attributable to the presence of mature pollen during the first experiment, while in the second experiment the anthers were at an immature stage prior to pollen release. Pollen grains have been suggested to serve as nutrient sources not only for fungi but also for bacteria (Scherman et al., 2025). In addition, we observed fungal growth from one spikelet to another, and that hyphae tended to colonize the anthers first, which might also represent a suitable niche for bacterial colonization. Therefore, the conditions of the first experiment likely facilitated enhanced bacterial growth and transport. In future studies, it would be valuable to determine whether the bacteria multiply during transport and/or only once they reach a new niche full of nutrients. In this way, the underlying cause(s) of the differences observed between the two *in planta* experiments can be formally evaluated. Furthermore, bacteria were detected not only beyond the inoculation site but also at the inoculation site after 5 dpi, indicating that the bacterial strains can survive *in planta* during fungal infection. Therefore, the observed pattern appears to reflect expansion rather than relocation similar to what was observed in the 3D-printed device.

*Escherichia coli* K-12 was included as a negative control due to its inability to disperse with *F. graminearum* in the 3D-printed device. However, we observed that *E. coli*-RFP can be transported by *F. graminearum* in both *in planta* experiments. Although *E. coli* transport was less efficient (4 out of 6 replicates) than that of *P. hospita* (6 out of 6 replicates), it would be interesting to determine whether *F. graminearum* mediated bacterial transport is bacterial species specific. Future studies in which spikelets below the inoculation site are individually processed would allow assessment of how strong the association is between each bacterial strain and the fungus, enabling clearer discrimination between strains with strong fungal associations and those with weaker ones. Overall, the *in planta* method developed in this study validates the *P. hospita* transport by *F. graminearum* that was observed in the 3D-printed device and demonstrates that the device successfully mimics key aspects of FH evident *in planta* in a natural host niche.

### Toward a multifactorial understanding of FH

Our results support a multifactorial model of bacterial movement along hyphae, shaped by both biotic and abiotic factors. While some fungi, such as *C. cinerea*, act as poor transporters overall, certain bacteria still reached the target well under specific conditions, emphasizing the combined influence of fungal traits, bacterial motility, and environmental context. For example, in rich medium, the oomycete *P. ultimum* transported *P. koreensis* VD-NE, whereas the fungus *F. graminearum* did not; conversely, *F. graminearum* transported *P. hospita* while *P. ultimum* failed.

Transport mechanisms may vary with inoculation strategy. In “cis” setups (co-inoculation), bacteria crossed as early as 3 dpi, suggesting potential passive transport via water flow associated to hyphal growth, whereas “trans” setups (opposite wells) likely required active motility. Interestingly, *P. putida* KT2440, despite its known motility, crossed less efficiently in “trans” conditions, indicating that bacterial energy allocation and environmental cues may influence dispersal. These findings align with previous studies showing that water films alone can enable movement as suggested by Ruan et al., (2022) and that distance reduces transport efficiency (Kuhn et al., 2022). Conversely, the efficient crossing in LMN of *F. cupreum*, the only supposedly non-motile strain tested, warrants further evaluation of the mechanisms of dispersal in this species.

Overall, there is growing understanding that FH are not universal conduits, but rather dynamic systems governed by mycelial growth patterns, bacterial strategies, medium composition, water availability, and distance. Our device provides a platform to disentangle these variables and advance a more nuanced understanding of BFI. Future work should integrate real-time imaging and controlled environmental gradients to clarify the interplay between passive and active transport mechanisms.

## Conclusion

We present a novel method with increasing throughput for the screening of potential FH partners. Our study highlights the interplay of both biotic and abiotic factors, which render difficult the prediction of interaction outcomes. Key insights on factors influencing bacterial translocation include medium composition, inoculation strategy, fungal exploratory behavior, and intra-strain variability of motility and potentially chemotactic behavior. Notably, we observed that even organisms deemed as "poor transporters" can enable bacterial dispersal under specific conditions, underscoring the context-dependent nature of these interactions. Likewise, non-motile bacteria have been able to disperse in some instance, indicating that passive disperal also plays a role. An intriguing observation made with the devices was the replacing of bacteria associated with fungal exploratory behavior. Those instances were the most numerous in nutrient-poor environments with a highly exploratory fungus, suggesting the existence of multiple FH strategies among mycelia-forming organisms in response to their environment. Together, our findings demonstrate highly variable outcomes in FH interactions. Our framework encourages future studies to adopt a layered experimental approach which could account for factors such as nutrient gradients, timing of inoculation, and volatile signaling to unravel the mechanisms underlying FH. Ultimately, we present our device as a new tool for better understanding of the rules governing FH formation and more systematic investigations of the chemical and physical cues that modulate bacterial motility and fungal exploration in structured environments.

## Acknowledgements

This research was supported by a Science Focus Area Grant from the U.S. Department of Energy (DOE), Biological and Environmental Research (BER), Biological System Science Division (BSSD) under grant number LANLF59T. MD and KHK contributions to this study were supported by BBSRC International Institutional Award Tranche 1 (BB/Y514196/1) and the BBSRC Delivering Sustainable Wheat Institute strategic programme (BB/X011003/1) and the constituent work packages (BBS/E/RH/230001B). AN and PK contributions to this study were supported by the U.S. National Science Foundation under grant number 2038293. The *Fusarium graminearum in planta* study was done under the conditions specified in the UK APHA licence101948/198285. We would like to thank Dr. Tim Mauchline and Dr. Susan Mosquito from the Microbiology Group at Rothamsted Research for kindly providing the pBBR-RFP vector and the bacterial transformation protocol.

## Supplementary material

**Supplementary Information 1:** .stl files for devices and holders.

**Supplementary Table 1.**
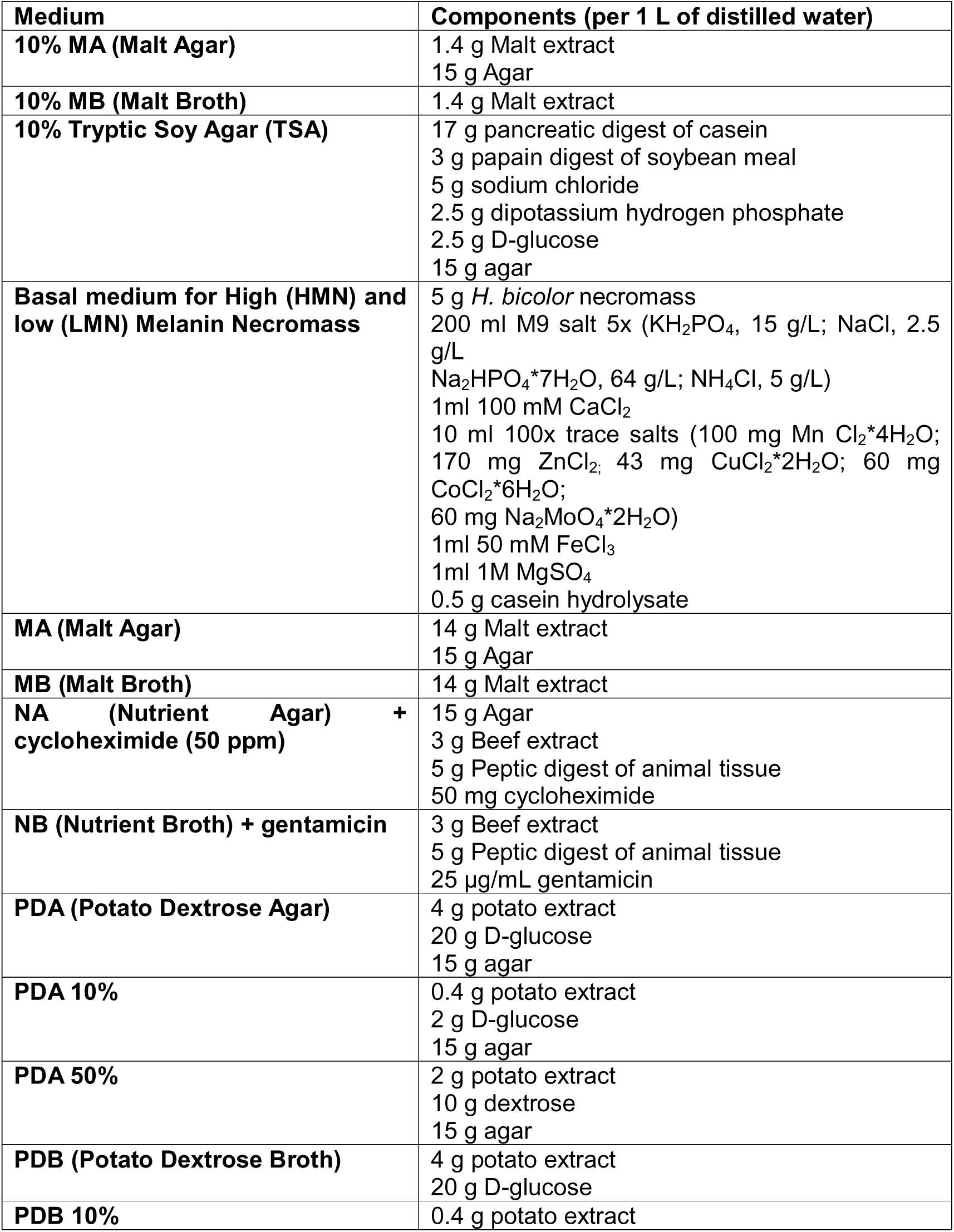
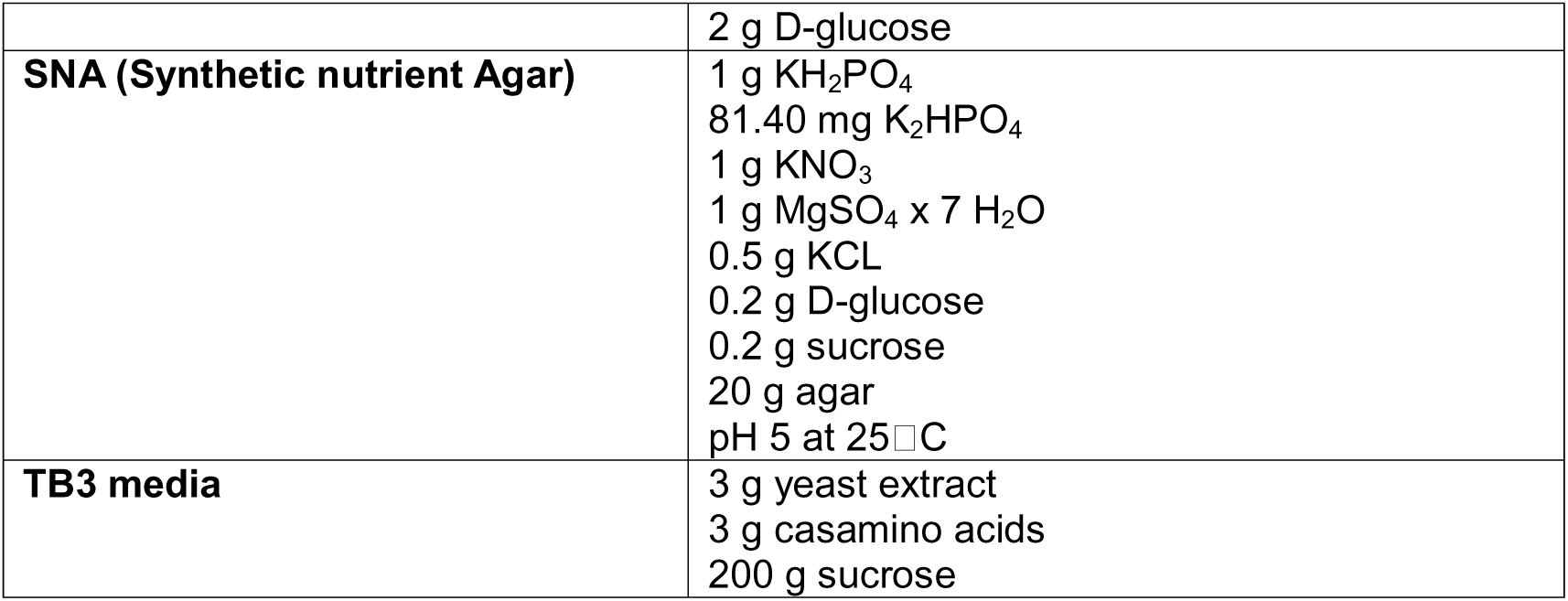
Media composition.

**Supplementary Figure 1.**
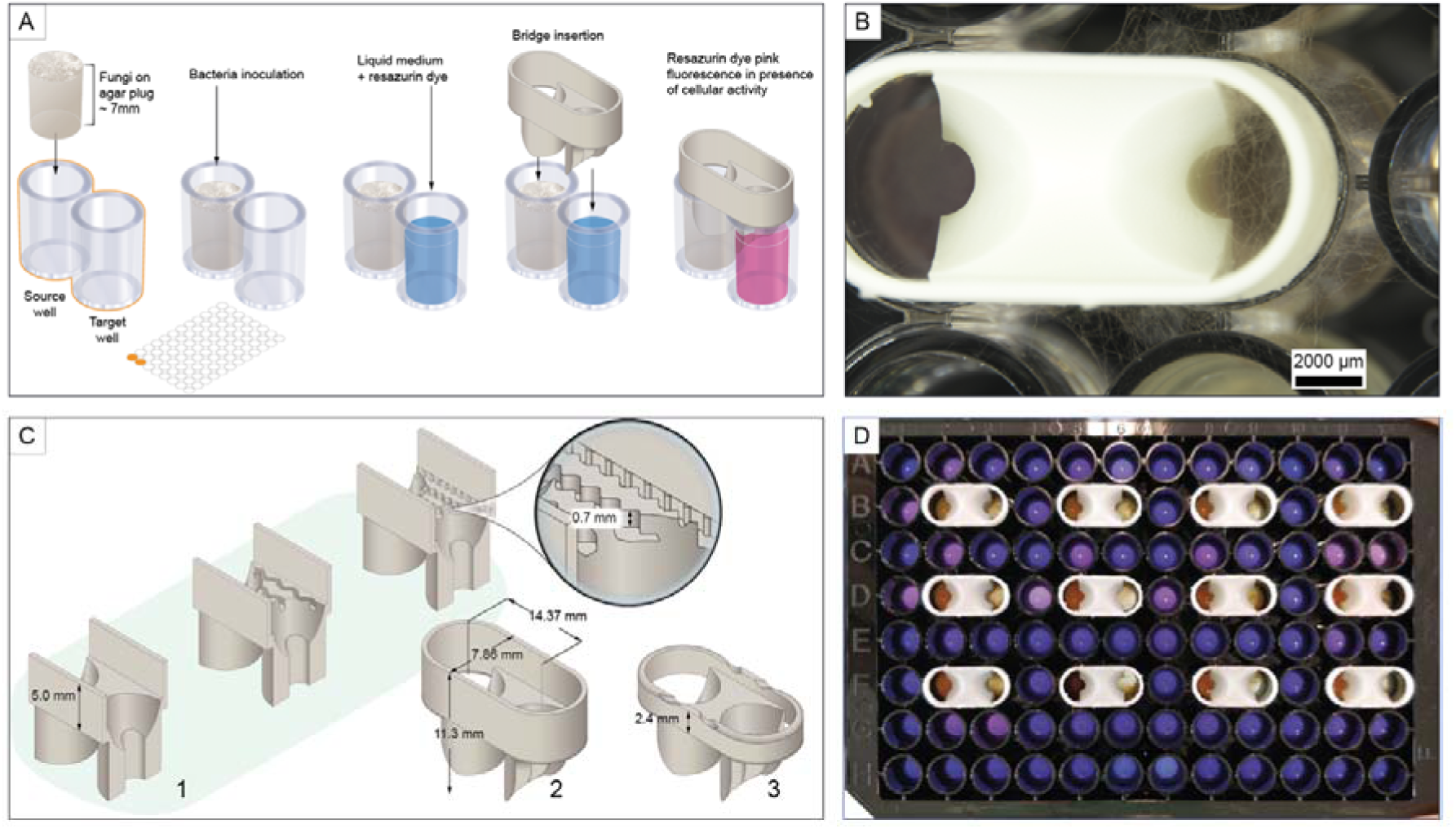
Initial design of the fungal highways bridge device. (**A**) The initial device was designed to be used as an add-on to a 96-well plate. Two adjacent wells acted as source and target wells. Each device was designed to leave enough space for the insertion of an agar plug for fungal inoculation in the source well. The addition of resazurin dye in liquid medium in the target well was intended to allow fast assessment of microbial development signaled by the color change of this metabolic dye. (**B**) Image of the inoculated device observed from the top. A major problem with this device was hyphal colonization of adjacent wells, leading to cross-contamination as observed in the target well. (**C**) Examples of iterative design validation. Lane 1 shows the different surface morphologies that were tested. Lanes 2 and 3 show examples of modification on the height of the enclosing wall. (**D**) Evaluation of the device model shown in panel C Lane 3 in a 96-well plate. The fungus was inoculated on the right side of the device. Uninoculated wells were filled with liquid medium containing resazurin to detect fungal colonization of adjacent wells as indicated by a change in dye color from violet to red.

**Supplementary Figure 2.**
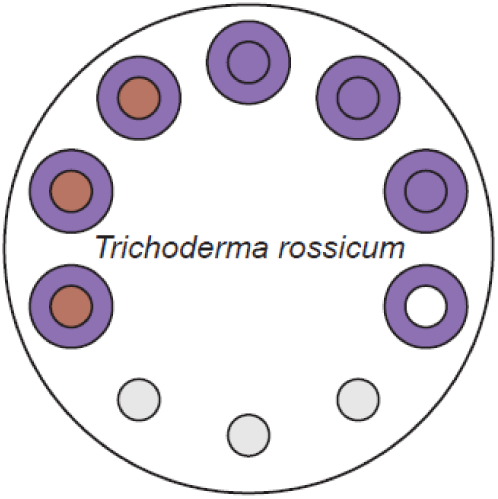
Performance assessment of the devices using fungal–bacterial pairings with two media types and the fungus *T. rossicum*. Layout of the wells and bacterial replacement in the target wells for 10% MB. The smaller circle indicates the expected outcome in the target medium, corresponding to the inoculated bacterial strains. The bigger (outer) circles indicate the actual outcome. As a result, same-colored circles indicate that expected and actual outcomes were the same.

**Supplementary Figure 3.**
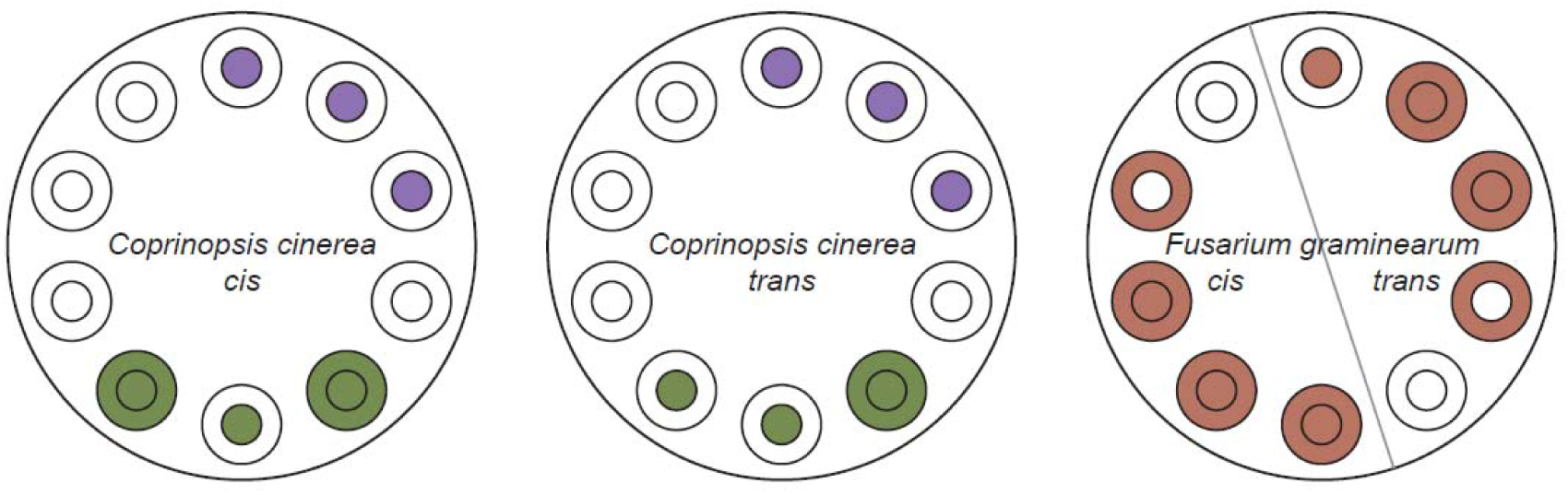
Assessing of FH interactions with two inoculation modes and different nutrient gradients. Sampling after 20 DPI. Layout of the wells and bacterial replacement in the target wells. The smaller circle indicates the expected outcome in the target medium, corresponding to the inoculated bacterial strains. The bigger (outer) circles indicate the actual outcome. As a result, same-colored circles indicate that expected and actual outcome were the same. Cis = co-inoculation in poor medium (10% MB), trans= inoculation of the fungus and the bacteria in separate wells.

**Supplementary Figure 4.**
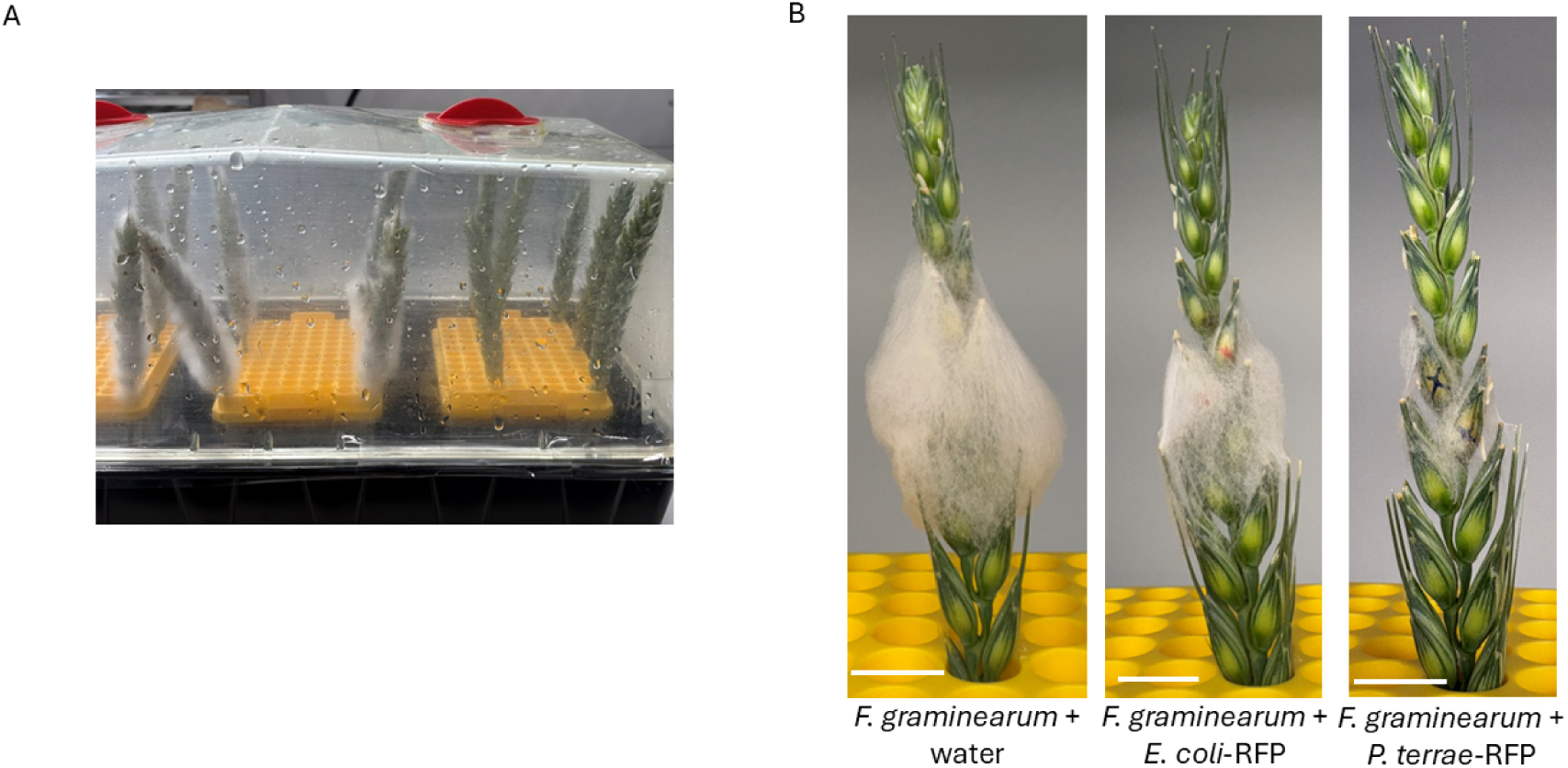
**(A)** Chamber used to incubate detached wheat spikes following inoculation. A wet mat was included to maintain high humidity, and tip boxes filled with water ensured continuous hydration of the spikes throughout the experiment. **(B)** Co-inoculation of *F. graminearum* with *P. hospita*-RFP reduces fungal growth. A mixture of *F. graminearum* spores with either *E. coli*-RFP or *P. hospita*-RFP was inoculated into two spikelets per wheat spike (noted by marks). Detached spikes remain hydrated and green throughout the experiment. However, the anthesis process is arrested in the basal, immature spikelets resulting in an overall arrest of spike maturation during the experimental period. Images were acquired at 5 dpi. Scale bar: 1 cm.

**Supplementary Figure 5.**
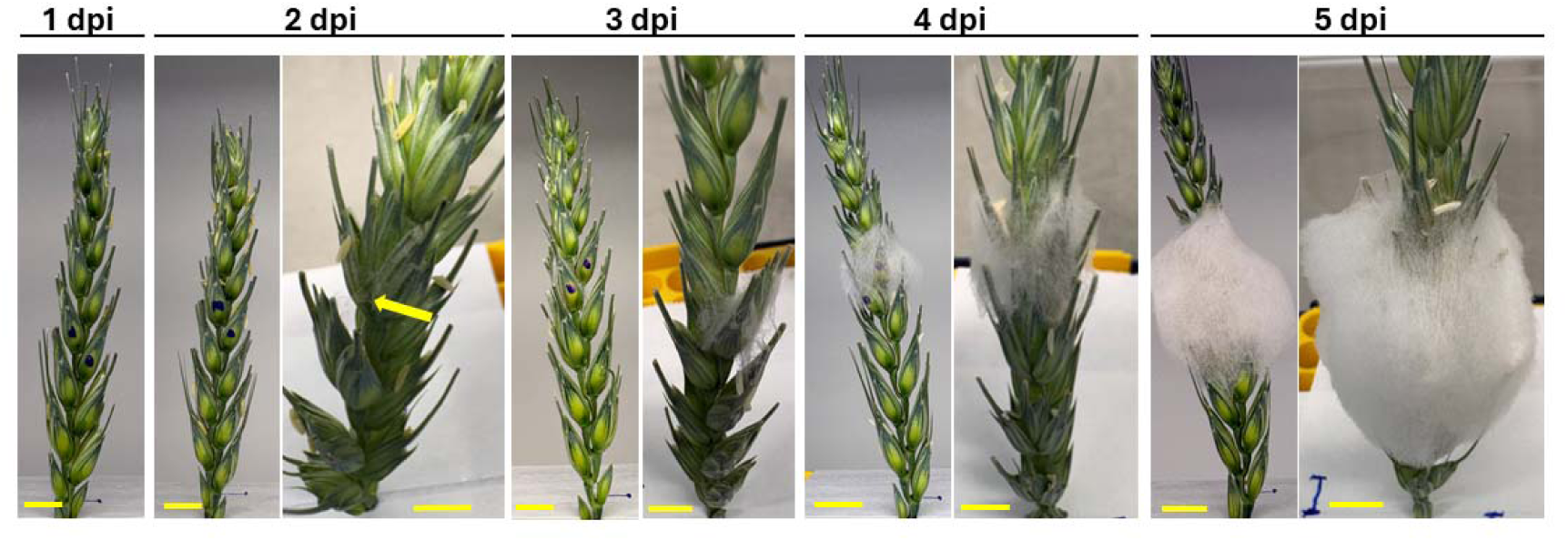
*Fusarium graminearum* growth during incubation under saturated humid conditions. Two spikelets from a wheat spike (marker with dots) were inoculated with *F. graminearum* spores. Inoculated spikes were detached from the plant and incubated in a chamber with saturated humid conditions. After 2 dpi (yellow arrow) mycelia can be observed outside the inoculated spikelets. Then, mycelia continue growing toward adjacent spikelets after 5 dpi. Scale bar: 1 cm.

**Supplementary Figure 6.**
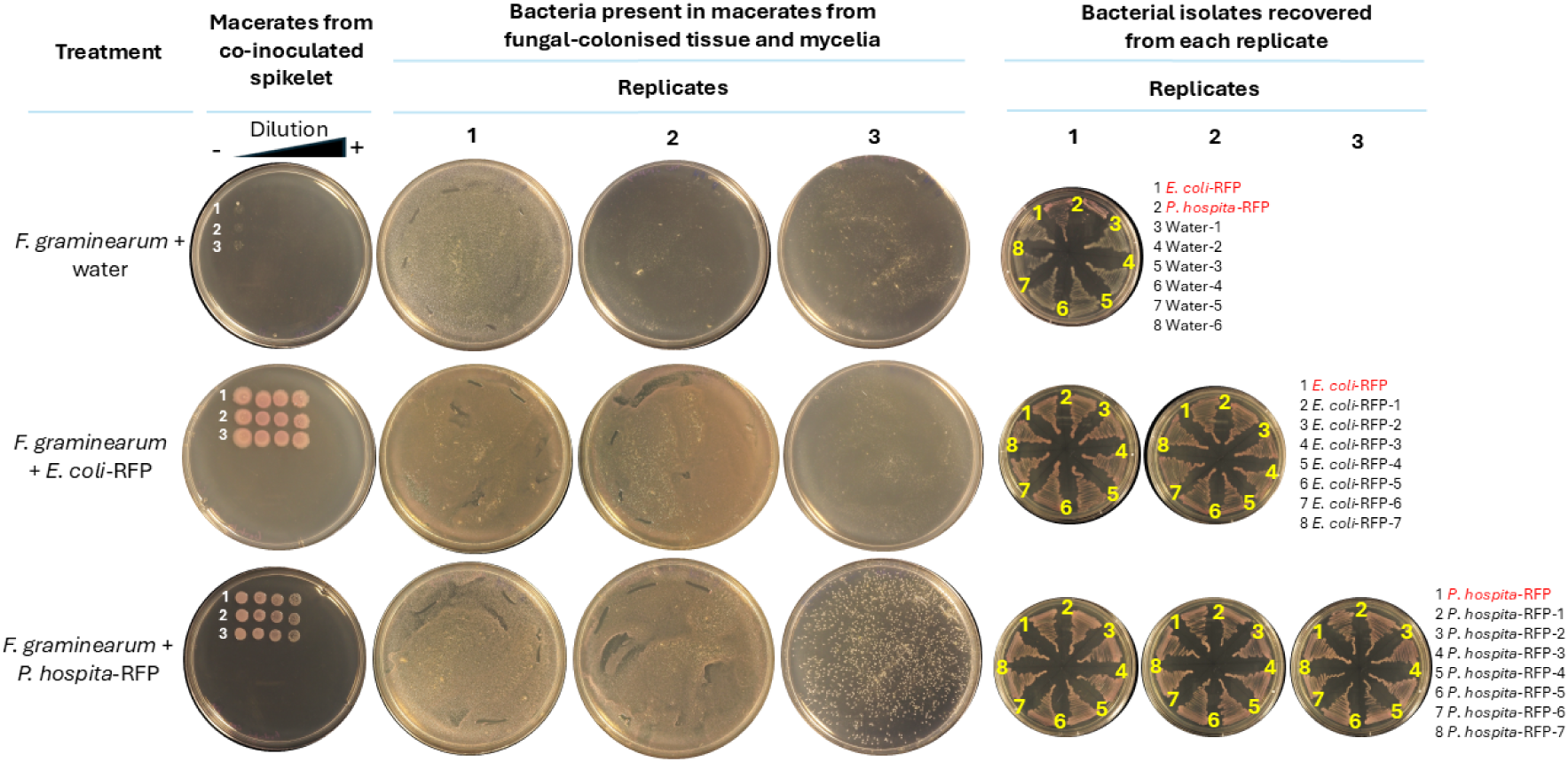
Bacteria recovered from macerates of tissue colonized by the fungus and mycelia. **Left panel**: Bacterial detection for each replicate (1 to 3). Co-inoculated spikelets for each treatment were macerated, serially diluted 10-fold, and plated on the same agar plate. Bacteria were detected in one replicate from *F. graminearum* inoculated with water (1), in 2 replicates from *F. graminearum* inoculated with *E. coli*-RFP (2 and 3), and in the 3 replicates from *F. graminearum* inoculated with *P. hospita*-RFP. **Right panel**: Bacteria recovered from each replicate. Colonies recovered from *E. coli*-RFP and *P. hospita*-RFP plates display a reddish coloration, as expected for these RFP-tagged strains. Numbered bacterial strains indicate bacteria recovered from the plates (shown in black) and control bacteria (shown in red).

**Supplementary Figure 7.**
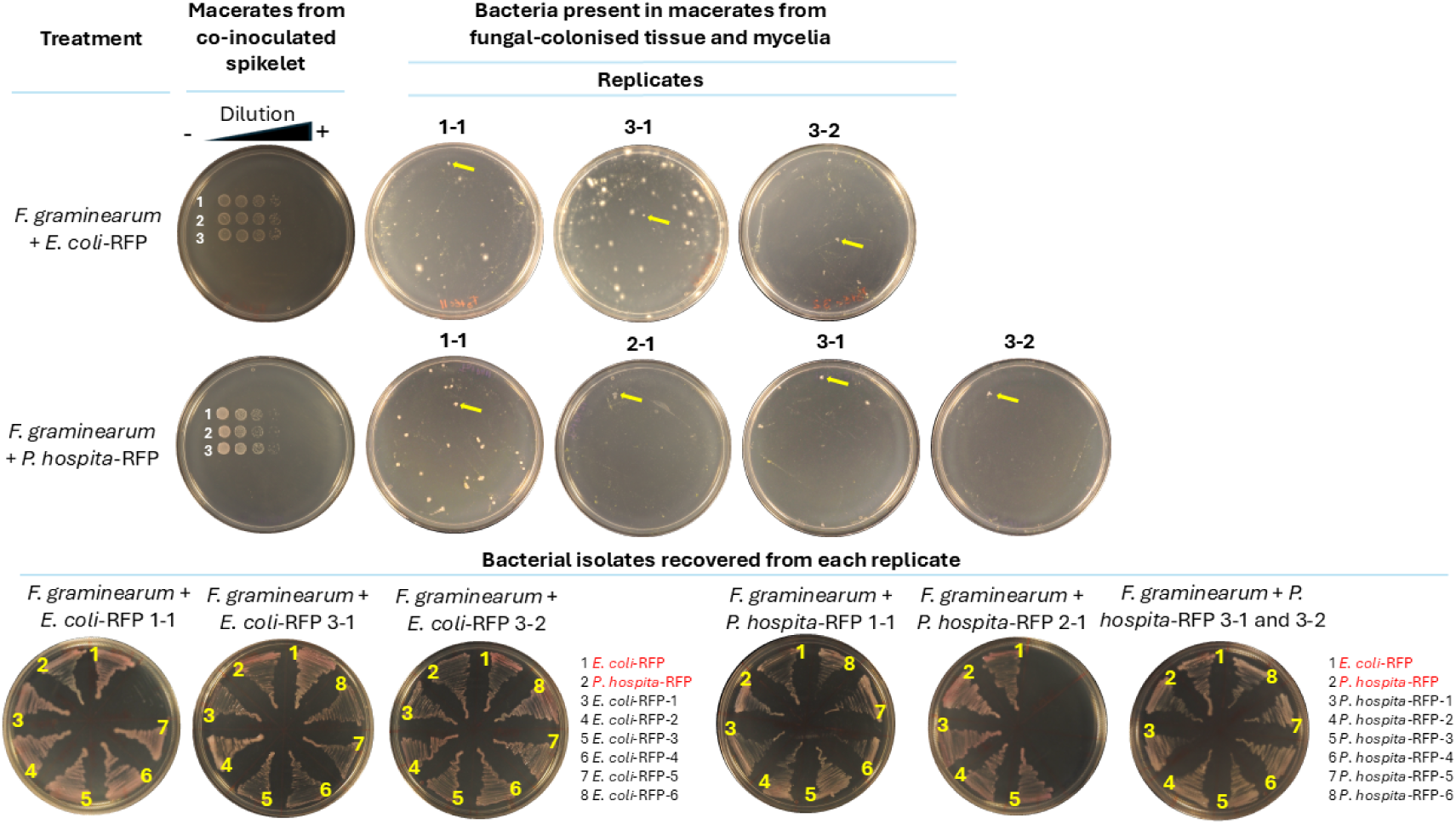
Bacteria recovered from macerates of tissue colonized by the fungus and mycelia. **Upper panel**: Detection of bacteria in macerates from spikes co-inoculated with *F. graminearum* and either *E. coli*-RFP or *P. hospita*-RFP. Co-inoculated spikelets for each treatment were macerated, serially diluted 10-fold, and plated on the same agar plate to confirm that the bacterial strains were present throughout the experiment. Bacteria were detected in two replicates for *F. graminearum* inoculated with *E. coli*-RFP (1-1 and 3-1), and in three replicates for *F. graminearum* inoculated with *P. hospita*-RFP (1-1, 2-1, and 3-1). Additionally, for both bacterial strains, bacteria were recovered from macerates of colonized spikelets from the spike positioned in front the co-inoculated spike (3-2). **Lower panel**: Bacteria recovered from each replicate. Colonies recovered from *E. coli*-RFP and *P. hospita*-RFP plates display the expected reddish coloration characteristic of RFP-tagged strains. Numbered bacterial strains indicate bacteria recovered from the plates (shown in black) and control bacteria (shown in red).

